# TIAM-1/GEF can shape somatosensory dendrites independently of its GEF activity by regulating F-actin localization

**DOI:** 10.1101/347567

**Authors:** Leo T.H. Tang, Carlos A. Díaz-Balzac, Maisha Rahman, Nelson J. Ramirez-Suarez, Yehuda Salzberg, María I. Lázaro-Peñal, Hannes E. Bülow

**Author notes:** contributed equally (Leo T.H. Tang, Carlos A. Díaz-Balzac). present address: Weizmann Institute, Rehovot, 7610001, Israel (Yehuda Salzberg).

## Abstract

Development of dendritic arbors is crucial for nervous system assembly, but the intracellular mechanisms that govern these processes remain incompletely understood. Here we show that the complex dendritic trees of PVD somatosensory neurons in *Caenorhabditis elegans* are patterned by distinct pathways downstream of the DMA-1 leucine rich transmembrane (LRR-TM) receptor. The guanine nucleotide exchange factor *tiam-1/GEF* and *act-4/Actin* function with the DMA-1/LRR-TM to pattern 4° higher order branches by localizing F-actin to the distal ends of developing dendrites. Biochemical experiments show that DMA-1/LRR-TM is part of a biochemical complex with TIAM-1/GEF and ACT-4/Actin. Surprisingly, TIAM-1/GEF appears to function independently of Rac1 guanine nucleotide exchange factor activity. Additionally, another pathway dependent on HPO-30/Claudin and TIAM-1/GEF is required for formation of 2° and 3° branches. Collectively, our experiments suggest that the DMA-1/LRR-TM receptor on PVD dendrites may control aspects of dendrite patterning by directly modulating F-actin dynamics, independently of TIAM-1/GEF enzymatic activity.

## Introduction

Neurons are highly polarized cells, which comprise a single axon and often elaborately sculpted dendritic arbors. Dendrites receive input from other neurons or the environment, whereas the single axon transmits information to other neurons. The nervous system is formed by a myriad of specific synaptic connections between neurons and the formation of these connections is influenced by the shape and complexity of dendritic arbors. Both genes that act within the developing neurons and in surrounding tissues are crucial to establish distict dendritic structures during development (JAN AND JAN 2010; DONG *et al.* 2015; LEFEBVRE *et al.* 2015). Of note, defects in dendrite morphology have been found in various neurological disorders (KAUFMANN AND MOSER 2000; KULKARNI AND FIRESTEIN 2012).

Both dendritic and axonal morphology is driven by the cytoskeleton and regulators of the cytoskeleton have consequently important functions in neuronal development. Major components of the cytoskeleton include actin and tubulin, which form filamentous polymers named F-actin and microtubules, respectively. F-actin exists in unbranched and branched forms, whereas microtubules are generally unbranched. These filament-like polymers are not static, but highly dynamic structures due to the constant association and dissociation of monomers at either end. A plethora of proteins bind to and modulate polymerization and depolymerization of both F-actin and microtubules in neurons (reviewed in (DENT *et al.* 2011; KAPITEIN AND HOOGENRAAD 2015; KONIETZNY *et al.* 2017)). F-actin and microtubules are important for countless aspects of neuronal function and development in both axons and dendrites, including differentiation, migration and the elaboration of axonal and dendritic processes (JAN AND JAN 2010; DENT *et al.* 2011).

Regulation of the cytoskeleton is controlled by dedicated signaling pathways, which often originate with cell surface receptors. These receptors utilize regulatory proteins such as guanine nucleotide exchange factors (GEFs) or GTPase activating proteins (GAPs), which, in turn, modulate the activity of small GTPases. For example, the RacGEF Tiam-1, first identified in flies (SONE *et al.* 1997), regulates activity-dependent dendrite morphogenesis in vertebrates (TOLIAS *et al.* 2005), likely by activating the small GTPase Rac1. Activated GTPases such as Rac1 then bind the WASP family verprolin-homologous protein (WVE-1/WAVE) regulatory complex (WRC) of actin regulators to promote actin polymerization and branching (CHEN *et al.* 2010).

The polymodal somatosensory neuron PVD in *Caenorhabditis elegans* has emerged as a paradigm to study dendrite development. The dendritic arbor of PVD neurons develops through successive orthogonal branching (OREN-SUISSA *et al.* 2010; SMITH *et al.* 2010; ALBEG *et al.* 2011)(Figure 1A). During the late larval L2 stage primary (1°) branches first emerge both anteriorly and posteriorly of the cell body along the lateral nerve cord. In subsequent larval stages secondary (2°) branches emanate orthogonally to bifurcate at the boundary between the lateral epidermis and muscle to form tertiary (3°) branches. These, in turn, form perpendicular quaternary (4°) branches to establish the candelabra-shaped dendritic arbors, which have also been called menorahs (OREN-SUISSA *et al.* 2010). Previous studies have shown that an adhesion complex consisting of MNR-1/Menorin and SAX-7/L1CAM functions from the skin together with the muscle-derived chemokine LECT-2/Chondromodulin II to pattern PVD dendrites. This adhesion complex binds to and functions through the DMA-1/LRR-TM leucine rich transmembrane receptor expressed in PVD neurons (LIU AND SHEN 2011; DONG *et al.* 2013; SALZBERG *et al.* 2013; DIAZ-BALZAC *et al.* 2016; ZOU *et al.* 2016). DMA-1/LRR-TM shows great similarity in domain architecture with the LRRTM family of leucine rich transmembrane receptors in humans (LauréN *et al.* 2003), but limited sequence homology (data not shown). The signaling mechanisms that operate downstream of the DMA-1/LRR-TM receptor in PVD dendrites have remained elusive.

**Figure 1.**
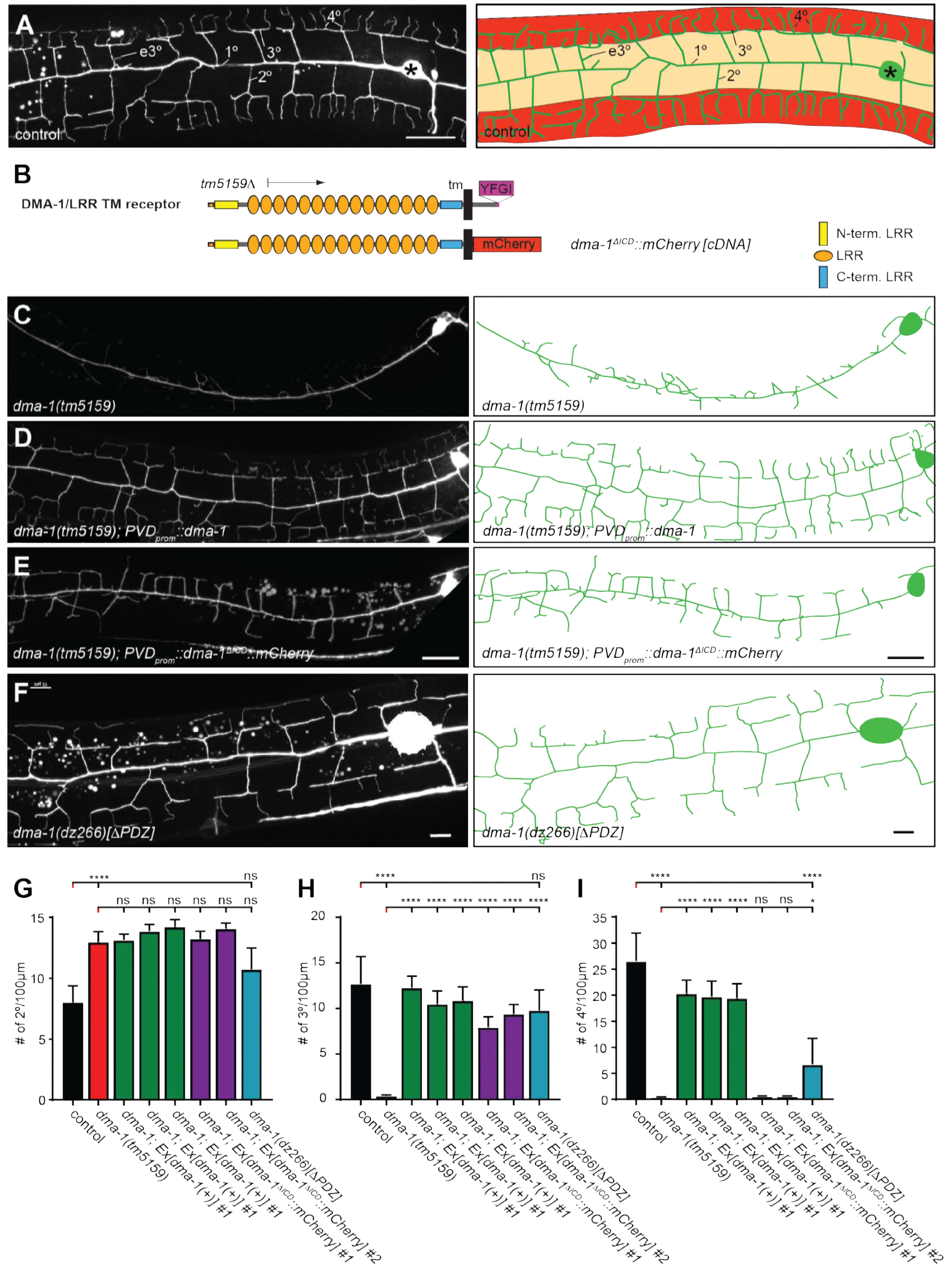
The intracellular domain of DMA-1/LRR-TM is required for higher order branching of PVD somatosensory dendrites. A. Fluorescent images of PVD (left panels) and schematics (right panels) of wild type control animals. PVD is visualized by the *wdIs52 [F49H12.4::GFP]* transgene in all panels. 1°, 2°, 3°, 4°, and ectopic 3° (e3°) dendrites are indicated. Anterior is to the left and dorsal is up in all panels, scale bars indicate 20μm. B. Schematics of the DMA-1/LRR-TM protein, including variants used in transgenic rescue experiments *(dma-1(ΔICD))* or predicted from CRISPR/Cas9 engineered knock-in alleles. A PDZ bindings site (YFGI) at the extreme C-terminus is indicated in lilac. The predicted deletion from the *tm5159* deletion allele is shown. C. - F. Fluorescent images of PVD (left panels) and schematics (right panels) of the genotypes indicated. Scale bar indicates 10μm. G. Quantification of 2°, 3°, and 4° branch numbers per 100μm anterior to the PVD cell body. Data for three and two independent transgenic lines for the *dma-1* wild type cDNA or the *dma-1(ΔICD)*, respectively, are shown next to the data for the *dma-1(dz244)* knock in allele. Data are represented as mean ± SEM. Statistical comparisons were performed using one-sided ANOVA with the Tukey correction. Statistical significance is indicated (****p < 0.0005). n = 10 animals per genotype.

Here we show that DMA-1/LRR-TM forms a complex with the claudin-like molecule HP0-30 (SMITH *et al.* 2013), which is required for localization of the DMA-1/LLR-TM receptor in PVD dendrites. Consistent with these observations, DMA-1/LRR-TM functions genetically in the menorin pathway together with *hpo-30/Claudin*, as well as the guanine nucleotide exchange factor (GEF) *tiam-1/GEF* and *act-4/Actin*. The signaling complex is required for both the correct localization of F-actin to, and the exclusion of microtubules from the distal endings of developing somatosensory dendrites. Intriguingly, TIAM-1/GEF functions independently of its Rac1 guanine nucleotide exchange factor activity. Biochemical experiments show that DMA-1/LRR-TM can form a complex with TIAM-1/GEF and ACT-4/Actin. Collectively, our experiments suggest that the DMA-1/LRR-TM receptor can modulate F-actin dynamics and localization through TIAM-1/GEF in developing dendrites independently of GEF activity.

## Results

### Isolation of genes that function in PVD and FLP dendrites

From genetic screens for mutants with defects in PVD patterning (see Suppl. Experimental Procedures for details), we obtained recessive alleles of the leucine rich repeat (LRR) single pass transmembrane (TM) receptor *dma-1/LRR-TM* and claudin-like *hpo-30* (Figure 2A-B, Figure S1A-D), both of which have been shown to be expressed and function in PVD for patterning of the dendritic arbor (LIU AND SHEN 2011; SMITH *et al.* 2013). In addition, we isolated mutant alleles in *tiam-1* (Figure 2C, Figure S1E), the *C. elegans* homolog (DEMARCO *et al.* 2012) of the multidomain vertebrate Rac1 guanine nucleotide exchange factor (GEF) Tiam1 (T-Lymphoma Invasion And Metastasis-Inducing Protein 1)(HABETS *et al.* 1994).

**Figure 2.**
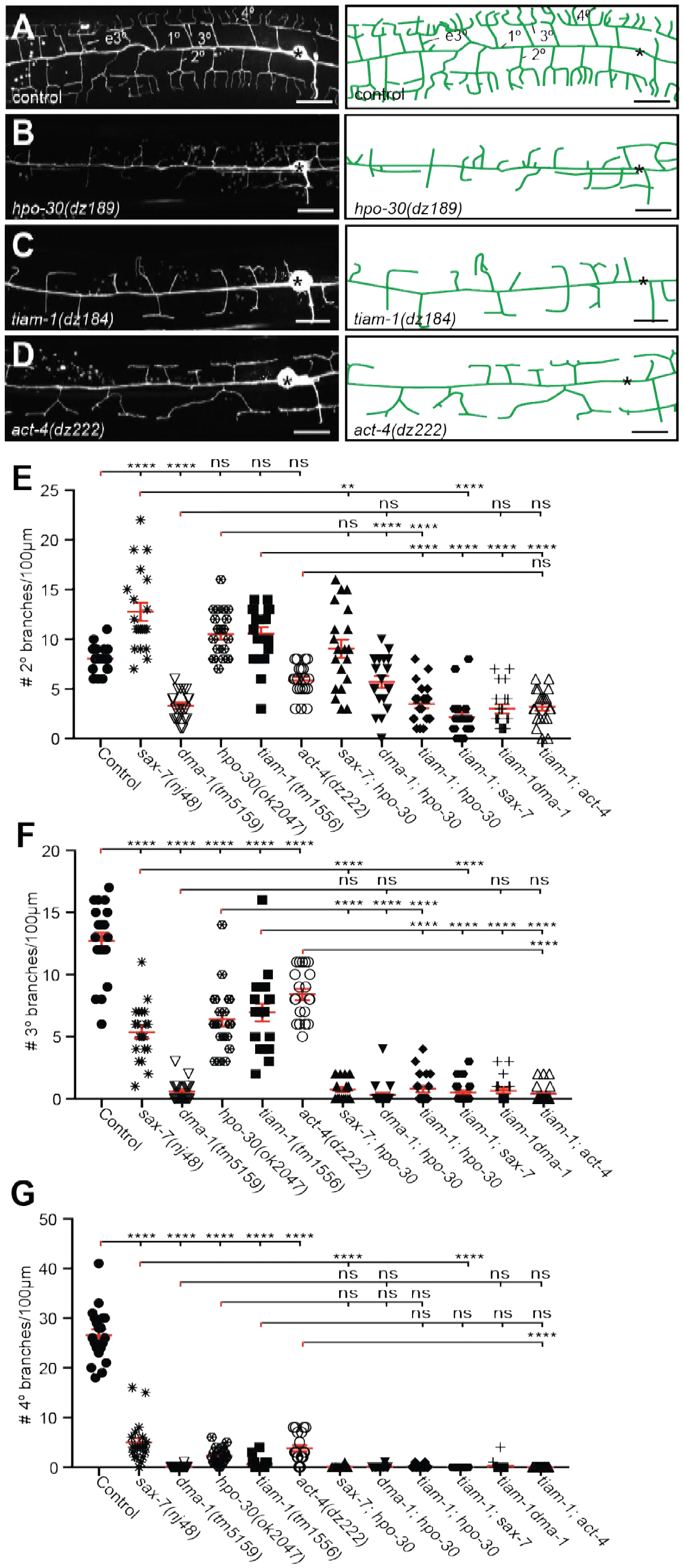
The *hpo-30/Claudin, tiam-1/GEF*, and *act-4/Actin* act genetically in the Menorin pathway. A. - D. Fluorescent images of PVD (left panels) and schematics (right panels) of the genotypes indicated. The control image is identical to Figure 1A and shown for comparison only. Details on alleles of individual genes and images of other alleles are shown in Figure S1. Scale bar indicates 20μm. E. - G. Quantification of 2°, 3°, and 4° branch numbers per 100μm anterior to the PVD cell body. Data for additional single and double mutants of the Menorin pathway as well as the average length and aggregate length of secondary, tertiary, and quaternary branches are shown in Figure S3. Data are represented as mean ± SEM. Statistical comparisons were performed using one-sided ANOVA with the Tukey correction. Statistical significance is indicated (****p < 0.0005). n = 20 animals for all samples.

We found that transgenic expression of *tiam-1/GEF* with heterologous promoters in PVD neurons but not in other tissues efficiently rescued *tiam-1/GEF* mutant phenotypes,consistent with expression of a *tiam-1/GEF* reporter in PVD neurons (DEMARCO *et al.* 2012)(Figure S1F-H). Finally, we isolated a mutant allele of *act-4/Actin* (Figure 2D, Figure S1I), one of five actins encoded in the *C. elegans* genome (KRAUSE *et al.* 1989). A reporter for *act-4* is expressed in muscle ((STONE AND SHAW 1993), data not shown), but fluorescent *in situ* hybridization experiments also suggested neuronal expression (BIRCHALL *et al.* 1995). We found that transgenic expression of *act-4/Actin* in PVD but not in muscle robustly rescued *act-4* mutant defects (Figure S1J-M). Additionally, expression of *act-1*, a paralog of *act-4*, rescued *act-4* mutant phenotypes (Figure S1M). All isolated mutant alleles also affected patterning of FLP neurons, a related pair of neurons, which cover the head region of the animal with similar dendritic arbors (Figure S2A). We conclude that in addition to DMA-1/LRR-TM and HP0-30/Claudin, TIAM-1/GEF and ACT-4/Actin function cell-autonomously to pattern the dendritic arbor of PVD and, likely FLP neurons. Moreover, expression of any actin in PVD rather than a specific function of ACT-4/Actin is important for dendrite patterning in PVD neurons.

### The PDZ binding site of the DMA-1 leucine rich transmembrane receptor (DMA-1/LRR-TM) is required for patterning of 4° branches but not 3° branches

Complete removal of *dma-1/LRR-TM* results in almost complete absence of 3° and 4° branches with additional effects on the number of 2° branches (LIU AND SHEN 2011; DONG *et al.* 2013; SALZBERG *et al.* 2013)(Figure 1B,C). As previously shown (LIU AND SHEN 2011), we found that the *dma-1* mutant phenotype can be fully rescued by transgenic expression of a wild type DMA-1/LRR-TM cDNA (Figure 1C,D,G). In contrast, a mutant where the intracellular domain (ICD) of DMA-1 was replaced by mCherry (ΔICD) resulted in partial rescue, where 3° branches, but not 4° branches were restored (Figure 1 B,C,E,G). We noticed that the C terminus of DMA-1/LRR-TM follows the consensus sequence of a PDZ binding site (X-Φ-X-Φ). We therefore used CRISPR/Cas9 genome editing to generate mutant animals lacking the last 4 residues of *dma-1* (referred to as ΔPDZ thereafter). The 1°, 2° and 3° branches of these mutants appear indistinguishable from wild type animals, whereas the number of 4° branches is significant decreased (Fig. 1G-I). These results indicate (1) that the intracellular domain of DMA-1/LRR-TM is required for the formation of PVD higher order branches, and (2) that the PDZ binding site at the extreme C-terminus of DMA-1/LRR-TM is important for formation of 4 ° branches.

### *The hpo-30/Claudin, tiam-1/GEF* and *act-4/Actin genes act in the menorin pathway*

To better understand the function of *dma-1/LRR-TM, hpo-30/Claudin, tiam-1/GEF* and *act-4/Actin* and test their genetic interactions with the Menorin pathway, we investigated the PVD mutant phenotype in single and double mutants using morphometric analyses as previously described (SALZBERG *et al.* 2013). We determined the number and aggregate length for all classes of dendrites in a segment 100μm anterior to the PVD cell body in different genetic backgrounds. We found the number of 2° branches unchanged in *hpo-30/Claudin, tiam-1/GEF* and *act-4/Actin* single mutants compared to wild type animals. Double mutants between *sax-7/L1CAM* and *hpo-30/Claudin* were not more severe than the more severe of the single mutants, indicating that *hpo-30/Claudin* functions in the menorin pathway for 2° branch patterning (Figure 2E). However, the *dma-1/LRR-TM; hpo-30/Claudin* and *dma-1/LRR-TM; tiam-1/GEF* or, the *tiam-1; act-4* double mutant were statistically indistinguishable from the *dma-1/LRR-TM* single mutant, suggesting that *dma-1/LRR-TM* is epistatic and required for most if not all functions during patterning of higher order branches in PVD. Interestingly, double mutants between *tiam-1/GEF* and *mnr-1/Menorin, lect-2/Chondromodulin II, kpc-1/Furin, sax-7/L1CAM*, or *hpo-30/Claudin* are more severe than either of the single mutants alone, but indistinguishable from the *dma-1/LRR-TM* single mutant (Figure 2E, Figure S3K). These findings suggest that *tiam-1/GEF* also serves in a genetic pathway that functions in parallel to *mnr-1/sax-7/lect-2/hpo-30*. Similar genetic relationships were observed regarding the number of 3° branches and the aggregate length of 2° and 3° branches with one notable exception. Double mutants between *sax-7/L1CAM* and *hpo-30/Claudin* displayed an enhanced phenotype for 3° branches that was statistically indistinguishable from the *dma-1/LRR-TM* single mutant phenotype (Figure 2F, Figure S3), suggesting a parallel function for HP0-30/Claudin for higher order branching. Common to all single and double mutants was the near complete absence of 4° branches (Figure 2G, Figure S3). Together, our results suggest that (1) *hpo-30/Claudin* and *tiam-1/GEF* act in the Menorin pathway to pattern PVD dendritic arbors and, that (2) *hpo-30/Claudin* and *tiam-1/GEF* may also serve functions independently of the menorin pathway.

We next sought to order *dma-1/LRR-TM, hpo-30/Claudin, tiam-1/GEF*, and *act-4/Actin* within the genetic pathway using a combination of gain and loss of function approaches. Previous work showed that expression of the hypodermally derived cell adhesion molecule *mnr-1/Menorin* in muscle of wild type animals results in the appearance of dendritic arbors that resembled African Baobab trees (Figure 3A)(SALZBERG *et al.* 2013). We used this *mnr-1/Menorin* gain of function *(gof)* phenotype in combination with loss of function mutations in *hpo-30/Claudin, tiam-1/GEF* and *act-4/Actin*. We found that loss of *hpo-30/Claudin, tiam-1/GEF* or *act-4/Actin* completely suppressed the formation of baobab-like dendritic arbors (Figure 3A-E) to the same extent as removal of other genes in the Menorin pathway such as *dma-1/LRR-TM, sax-7/L1CAM*, or *lect-2/Chondromodulin II* (SALZBERG *et al.* 2013; DIAZ-BALZAC *et al.* 2016). Moreover, protein localization of a LECT-2/Chondromodulin II or SAX-7/L1CAM reporter is not affected in *hpo-30/Claudin, tiam-1/GEF* or *act-4/Actin* mutants (Figure S2B,C). These findings suggest that *hpo-30/Claudin, tiam-1/GEF* and *act-4/Actin*, just like *dma-1/LRR-TM, sax-7/L1CAM*, or *lect-2/Chondromodulin II* function genetically downstream of *mnr-1/Menorin*.

**Figure 3.**
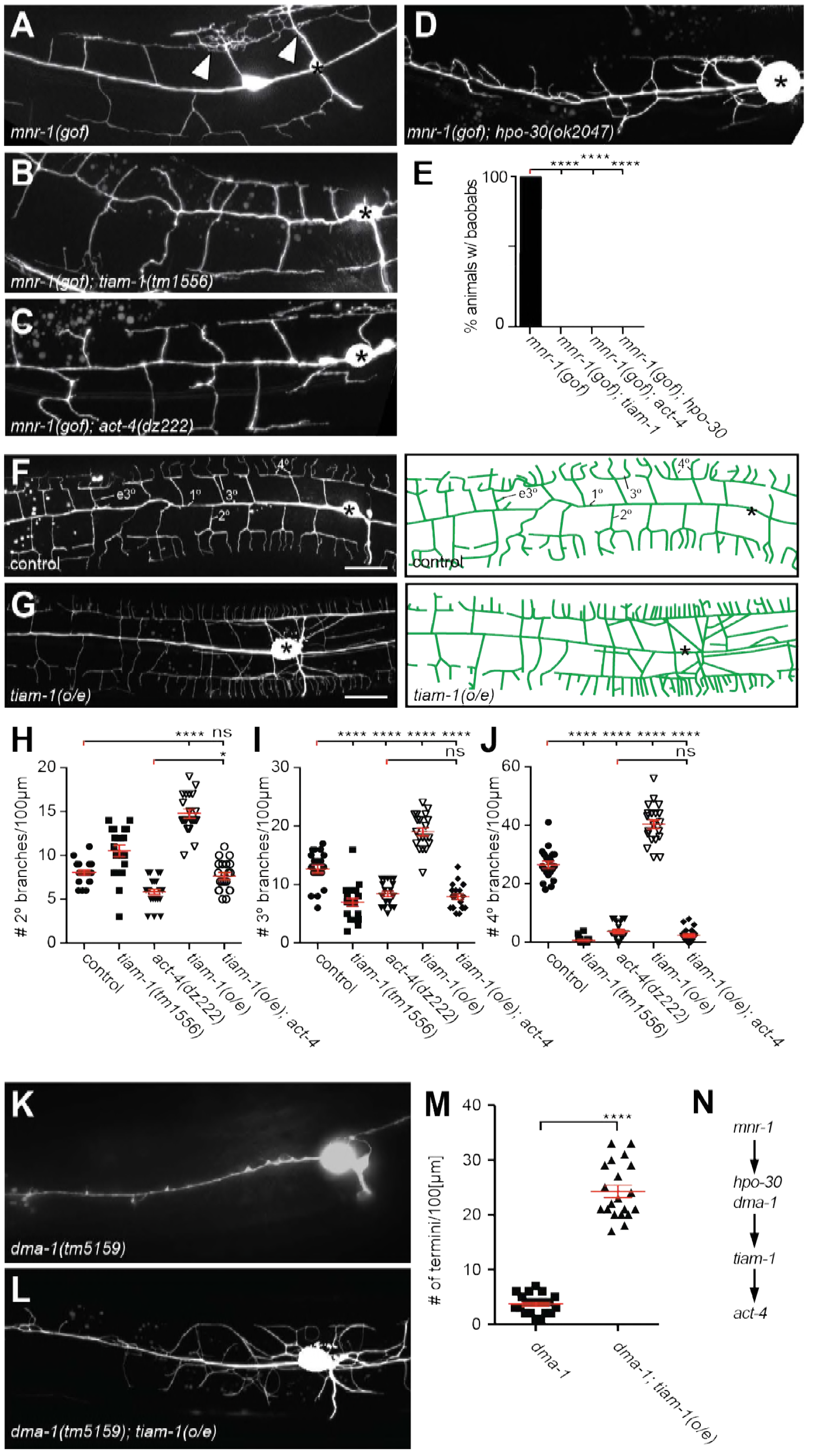
The *tiam-1/GEF* and *act-4/Actin* act downstream of the *dma-1/LRR-TM* receptor in PVD dendrites. A. - D. Fluorescent images of animals in which *mnr-1/Menorin* was expressed ectopically in muscle (*mnr-1(gof): dzIs43, Is[Pmyo-3::mnr-1]* (SALZBERG *et al.* 2013)) in different genetic backgrounds. A characteristic baobab-like tree is indicated by a white arrowhead in (A). PVD is visualized by the *wdIs52 [PF49H12.4::GFP]* transgene. Anterior is to the left, dorsal up, and scale bars indicate 20μm in all panels. E. Quantification of animals with baobab-like dendritic trees in the genotypes indicated. Data are represented as mean. Statistical comparisons were performed using the Z-test. Statistical significance is indicated (****p < 0.0005). n = 20 animals for all samples. F. - G. Fluorescent images of PVD (left panels) and schematics (right panels) of wild type control and *tiam-1(o/e)* overexpressing animals *(dzIs95, Is[Pser-2prom3::tiam-1])*. The control image (E) is identical to Figure 1A and shown for comparison only. H. - J. Quantification of secondary (2°, G), tertiary (3°, H), and quaternary (4°, I) branch numbers per 100μm anterior to the PVD cell body in the genotypes indicated. Data for control, *tiam-1(tm1556)*, and *act-4(dz222)* are identical to data in Figure 2 and shown for comparison only. Data are represented as mean ± SEM. Statistical comparisons were performed using one-sided ANOVA with the Tukey correction. Statistical significance is indicated (****p < 0.0005). n = 20 for all samples. K. - L. Fluorescent images of PVD in *dma-1/LRR-TM* mutant animals alone (J) and in combination with a *tiam-1(o/e)* expression array (K). Anterior is to the left and ventral down. Scale bar indicates 20μm. M. Quantification of dendrite termini in a 100μm section anterior to the PVD cell body in the genotype indicated. Data are represented as mean ± SEM. Statistical comparisons were performed using Student’s T-test. Statistical significance is indicated (****p < 0.0005). n = 20 animals for all samples. N. Putative genetic pathway from *mnr-1* to *act-4.*

We next asked in which order *dma-1/LRR-TM, tiam-1/GEF* and *act-4/Actin* function in PVD dendrites. Overexpression of *tiam-1/GEF (tiam-1(o/e))* in an otherwise wild type background resulted in an increase of 2°, 3°, and 4° branches in PVD dendrites (Figure 3F,G). This excessive branching was completely suppressed by a mutation in *act-4/Actin.* Specifically, the *tiam-1(o/e); act-4* double mutant was statistically indistinguishable from the *act-4* single mutant, both with regard to the number and aggregate length of dendritic branches (Figure 3H-J, Figure S4A-C). These observations suggest that *tiam-1/GEF* function is dependent on ACT-4/Actin in PVD dendrites. We further hypothesized that *tiam-1/GEF* would function downstream of the *dma-1/LRR-TM* receptor. If that were the case, one would predict that loss of branching in *dma-1/LRR-TM* null mutants would be at least partially suppressed (i.e. reversed) by overexpression of *tiam-1/GEF*. Indeed, overexpression of *tiam-1(o/e)* in *dma-1/LRR-TM* null mutants significantly increased the number of branches in PVD dendrites (Figure 3K-M). Taken together, our experiments reveal a pathway, in which TIAM-1/GEF functions downstream of or in parallel to DMA-1/LRR-TM (and by inference *hpo-30/Claudin*) at the PVD cell membrane and in a manner that is dependent on ACT-4/Actin (Figure 3N).

### HPO-30/Claudin functions to localize DMA-1/LRR-TM

Both DMA-1/LRR-TM and HP0-30/Claudin are predicted to be transmembrane proteins functioning in PVD dendrites. To determine the mechanistic relationship between these two factors, we asked where a functional DMA-1::GFP and a HPO-30::wrmScarlet protein fusion would be localized within the PVD neurons. As previously reported (DONG *et al.* 2016), the functional DMA::GFP reporter is localized in a punctate fashion in the cell body and along the dendritic tree as well as on the membrane (Figure 4A, upper panel). We found that a HP0-30::wrmScarlet fusion was also predominantly localized in a punctate fashion along the dendritic processes with less staining in the cell body or axon of PVD neurons (Figure 4A, middle panel), much like recently described HP0-30::GFP reporters (SMITH *et al.* 2013; ZOU *et al.* 2015). The DMA-1::GFP and HP0-30::wrmScarlet reporters show significant colocalization in dendritic processes, suggesting that they may be closely associated (Figure 4A, lower panel).

**Figure 4.**
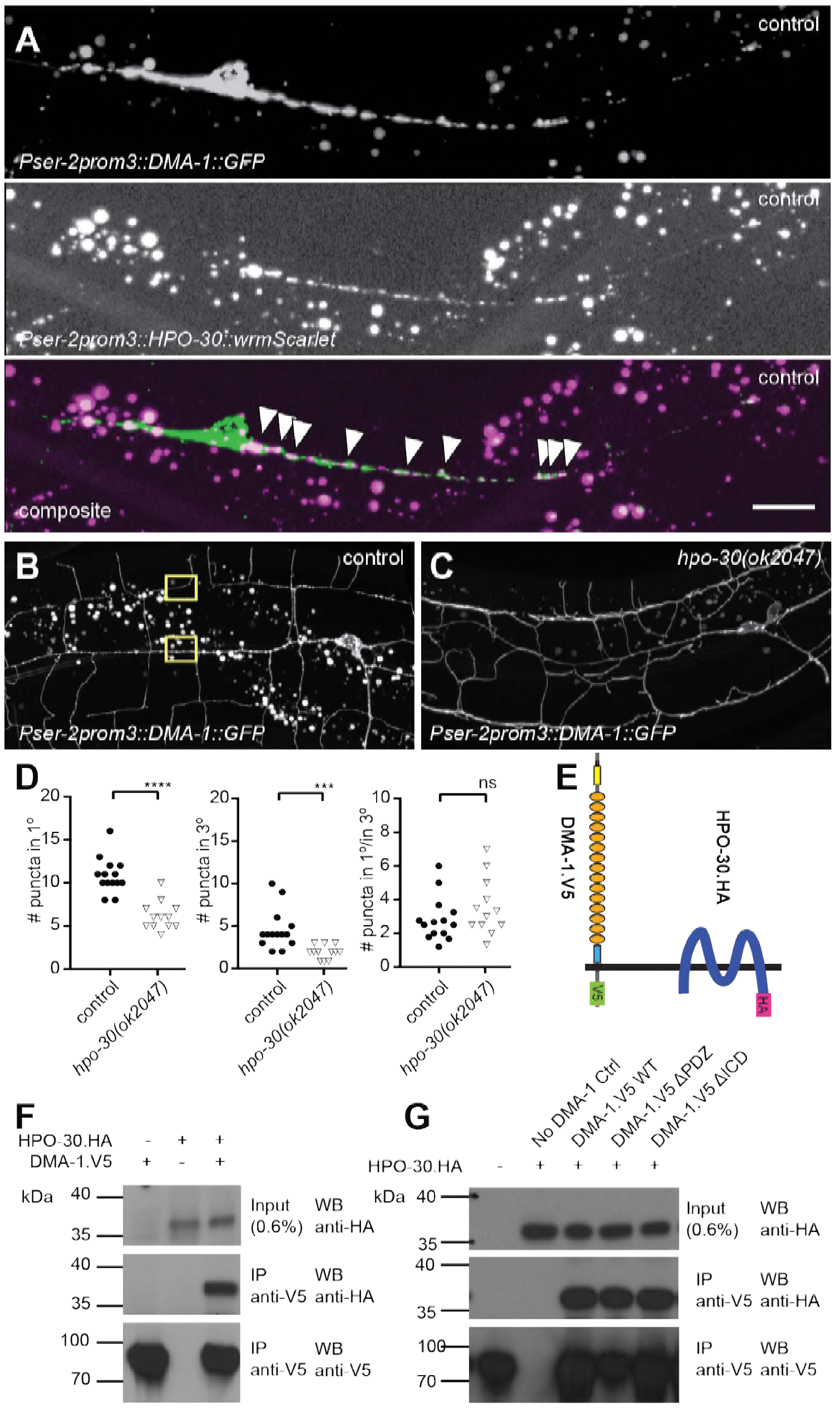
HPO-30/Claudin forms a complex with, and, localizes DMA-1/LRR-TM to higher order branches of PVD somatosensory neurons. A. Fluorescent images of animals expressing a functional DMA-1::GFP fusion (upper panel), a HP0-30::wrmScarlet fusion (middle panel), a merged image (lower panel). (wrmScarlet is a new, bright red fluorescing protein (EL MOURIDI *et al.* 2017)). Scale bars indicate 10 μm in all panels. White arrowheads indicate areas of colocalization. B. - C. Fluorescent images of PVD in animals expressing a functional DMA-1::GFP fusion in a wild type control (B) and a *hpo-30* mutant (C) background. Note the diffuse staining along the dendrite in *hpo-30* mutant animals compared to the punctate staining in a wild type control background. Yellow boxes indicate an area with puncta in 1° versus 3° dendrites. Puncta were quantified 30μm anterior to the cell body. D. Quantification of DMA-1::GFP puncta in the indicated genotype. n=14 wild type controls; n=12 *hpo-30(ok2047)* mutant animals. The number of DMA-1::GFP puncta in *tiam-1* and *act-4* mutants was also reduced in 1° dendrites, with variable effects on 3° dendrites (Figure S2E,D). Statistical comparisons were performed using Kruskal-Wallis-test. Statistical significance is indicated (ns: not significant, ***p<0.005, ****p < 0.0005). E. Schematic showing the topography of the DMA-1/LRR-TM single pass transmembrane receptor and the four transmembrane, claudin-like, HP0-30 protein. Immuno tags (V5 and HA) used for co-immunoprecipitation experiments are shown. F. Western Blots of co-immunoprecipitation experiments. Transfected constructs are indicated above the panels. Antibodies used for immunopreciptation (IP) and Western Blotting (WB) are indicated. A molecular marker is denoted on the left. Note, that the two lower panels are from a single Western blot, which was developed repeatedly with two different antibodies after stripping. G. Western Blots of co-immunoprecipitation experiments. Transfected constructs are indicated above the panels. DMA-1.HA ΔPDZ and DMA-1.HA ΔICD are lacking the PDZ binding site or the intracellular domain, respectively. Antibodies used for immunopreciptation (IP) and Western Blotting (WB) are indicated. A molecular marker is denoted on the left. Note, that the two lower panels are from a single Western blot, which was developed repeatedly with two different antibodies after stripping.

We next asked whether DMA-1::GFP localization was dependent on *hpo-30/Claudin*. We found that loss of *hpo-30/Claudin* resulted in more diffuse localization of DMA-1::GFP throughout the whole dendrite, with less punctate staining characteristic for DMA-1::GFP in wild type animals (Figure 4B-D). Moreover, the number of puncta in 1° and 3° branches was reduced in *hpo-30* null mutants, although their relative distribution remained unchanged (Figure 4D). These findings suggest that HP0-30/Claudin is required for correct trafficking or localization of DMA-1/LRR-TM, possibly at the membrane.

To determine whether DMA-1/LRR-TM and HP0-30/Claudin (Figure 4E) are part of the same biochemical complex, we conducted co-immunoprecipitation experiments. We transfected human embryonic kidney cells (HEK293T) with DMA-1 (tagged with HA, DMA-1.HA) and HP0-30 (tagged with V5, HP0-30.V5) alone and in combination. We found that DMA-1.HA efficiently co-precipitated HP0-30.V5 from a cellular lysate, suggesting that both proteins are part of the same biochemical complex (Figure 4F). This biochemical interaction was independent of the intracellular domain of DMA-1 (Figure 4G), suggesting that the proteins interact through either the transmembrane segments or the extracellular domains of DMA-1. Taken together our results are consistent with a model in which (1) HP0-30/Claudin exists in a complex with DMA-1/LRR-TM and (2) may serve a role in either correct trafficking or localization of DMA-1, or both within the dendritic arbor of PVD neurons, although we can not exclude additional functions for HP0-30 in signaling. It is interesting to note that HP0-30/Claudin has a similar membrane topology as stargazin, another claudin-like protein. Stargazin is required for synaptic targeting of ionotropic glutamate receptors (CHEN *et al.* 2000), possibly suggesting a more general function for claudin-like molecules besides their known role in the formation of tight junctions.

### The TIAM-1/GEF guanine nucleotide exchange factor functions independently of its Rac1 GEF activity to pattern PVD dendrites

To investigate the functions downstream of DMA-1/HP0-30 and to better understand putative downstream signaling events, we focused next on the function of the guanine nucleotide exchange factor TIAM-1. *C. elegans* TIAM-1/GEF is a multidomain protein (Figure 5A), which comprises, in order, a myristoylation signal, an EVH1 (Ena/Vasp homology) domain, a PDZ domain, and a DH/PH (Dbl homology domain/plekstrin homology domain) encoding the enzymatic guanine nucleotide exchange activity (DEMARCO *et al.* 2012). As suggested by work in flies and vertebrate neurons (SONE *et al.* 1997; KUNDA *et al.* 2001; TOLIAS *et al.* 2005), *C. elegans* TIAM-1/GEF has been shown to shape neurons through mechanisms dependent on the small GTPases *mig-2/RhoG* and *ced-10/Rac1* (DEMARCO *et al.* 2012). In addition to *mig-2/RhoG* and *ced-10/Rac1*, the *C. elegans* genome encodes one additional Rac1-like GTPase, *rac-2* (LUNDQUIST *et al.* 2001). Surprisingly, neither mutations in *mig-2/RhoG* and *ced-10/Rac1* alone or in combination, nor a mutation in *rac-2* resulted in obvious phenotypes in PVD dendritic arbor formation (Figure S5A-C), implying that TIAM-1/GEF functions independently of these Rac1-like GTPases, at least individually. To further investigate this notion, we conducted transgenic rescue experiments with deletions and point mutant variants of TIAM-1/GEF. We found that full length TIAM-1/GEF alone or as a C-terminal fusion with mCherry fully rescued the PVD dendrite patterning defects (Figure 5A, Figure S5D). Surprisingly, a T548F point mutant of full length TIAM-1/GEF still rescued the PVD mutant phenotype (Figure 5B, Figure S5D). The fact that the very same mutation in the *C. elegans* TIAM-1 DH/PH domain (DEMARCO *et al.* 2012) as well as the analogous point mutation in the UNC-73/Trio GEF abolishes Rac1 GEF activity *in vitro* (STEVEN *et al.* 1998), lent further support to the idea that TIAM-1/GEF activity may not be required for PVD patterning (Figure 5B-D). It was conceivable, however, that mutant TIAM-1(T548F) retained some residual GEF activity that through high level transgenic expression could provide sufficient enzymatic GEF activity to rescue the PVD dendrite defects of *tiam-1* mutants. To exclude this possibility, we used CRISPR/Cas9 genome editing to engineer the T548F missense mutation into the endogenous *tiam-1/GEF* locus. Two independently isolated alleles (*dz264* and *dz265*) that harbored the T548F point mutation in the *tiam-1/GEF* locus appeared phenotypically normal in regard to PVD patterning (Figure 5C-E and data not shown). Thus, a point mutant form of TIAM-1 (T548F) devoid of measurable Rac1 GEF activity *in vitro* is sufficient for PVD patterning.

**Figure 5.**
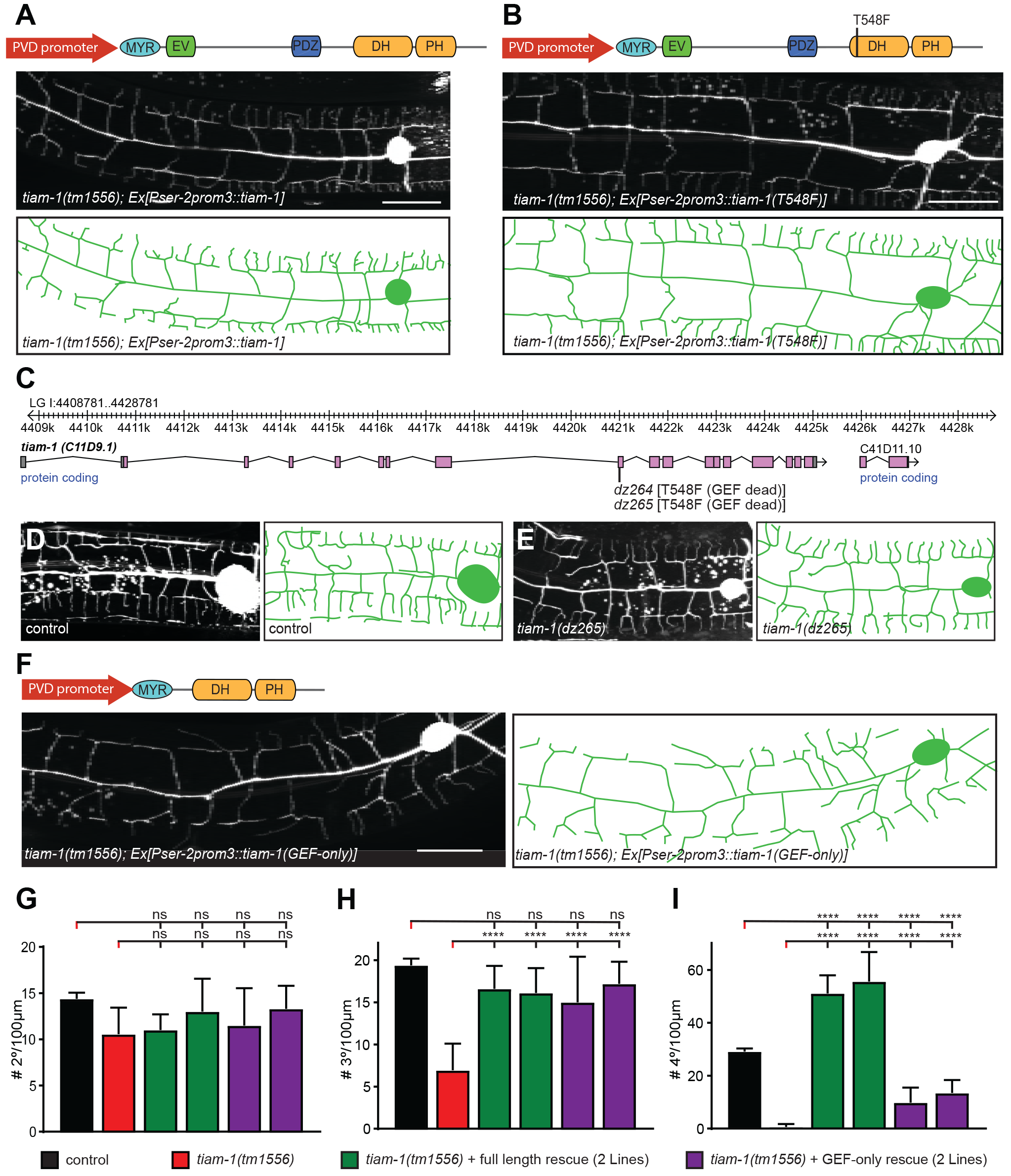
The TIAM-1/GEF functions independently of its guanine nucleotide activity. A. - B. Fluorescent images with schematics of *tiam-1* mutant animals carrying a wild type (A) or T548F point mutant form (B) of the *tiam-1* cDNA under control of the PVD specific *ser-2prom3* promoter. The constructs used in transgenic rescue experiment show schematically the TIAM-1/GEF multidomain protein, comprising in order a myristoylation signal (MYR), an enabled/vasodilator-stimulated phosphoprotein (VASP) homology domain (EV), a PDZ domain (PDZ) and, a Dbl homology (DH)/Plekstrin homology (PH) domain. The T548F point mutant (in B) is analogous to the S1216F mutation in UNC-73 (STEVEN *et al.* 1998) and the T516F mutation in a previously described TIAM-1 cDNA (DEMARCO *et al.* 2012). Importantly, this T516F point mutant has been shown to abrogate GEF activity towards Rac *in vitro* (DEMARCO *et al.* 2012). Scale bars indicate 20μm in all panels. C. Genomic environs of the *tiam-1* locus in linkage group I (LGI) with exons and introns indicated. The location of the two CRISPR/Cas9 engineered *dz264* and *dz265* alleles, encoding a missense mutation that effects the T548F mutation is shown. D. - E. Fluorescent images of PVD (left panels) and schematics (right panels) of wild type control (D) and *tiam-1(dz264)* mutant animals (E). PVD is visualized by the *wdIs52 [PF49H12.4::GFP]* transgene in all panels. F. Fluorescent image with schematics of *tiam-1* mutant animals carrying a deletion construct (GEF-only) of the *tiam-1* cDNA under control of the PVD specific *ser-2prom3* promoter. G. Quantification of 2°, 3°, and 4° branch numbers per 100μm anterior to the PVD cell body in the genotypes indicated. Data are represented as mean ± SEM. Statistical comparisons were performed using one-sided ANOVA with the Tukey correction. Statistical significance is indicated (****p < 0.0005). n = 10 animals per genotype.

Since the PDZ ligand motif of DMA-1/LLR-TM is important for the formation of 4° branches (Figure 1), we tested if the PDZ domain of TIAM-1/GEF may mediate this function. We found that truncated TIAM-1, lacking the EV and PDZ domains (referred to as GEF-only), only partially rescued the PVD mutant phenotype (Figure 5D, Figure S5). Specifically, the GEF-only construct rescued formation of PVD 2° and 3° defects in the same manner as the full-length TIAM-1 construct, but failed to rescue defects in the number of 4° branches (Figure 5E-G). Intriguingly, overexpression of full length TIAM-1 resulted in more 4° branches, suggesting that TIAM-1 function is dosage dependent (Figure 5G). Collectively, our results suggest that TIAM-1/GEF can function independently of its GEF activity and thus in a possibly GTPase-independent manner. Moreover, the DH/PH domain of TIAM-1 is sufficient for 2° and 3° branch formation, but not for 4° branch formation. These findings imply a role for the PDZ motif in DMA-1/LRR-TM and the PDZ binding domain in TIAM-1/GEF in mediating PVD 4° branch formation.

### F-Actin is localized to the leading edge of dendrites

The filamentous F-actin and microtubule polymers are part of the cytoskeleton that provide stability and force during growth and development of neuronal processes (DENT *et al.* 2011; KAPITEIN AND HOOGENRAAD 2015). Having identified a function for *act-4/Actin* in PVD dendrite patterning, we sought to visualize F-actin in PVD neurons of live animals. To this end, we used the calponin homology domain of utrophin (UtrCH) fused to tagRFP (tagRFP::UtrCH), which has been previously shown to faithfully visualize F-actin without effects on actin dynamics (BURKEL *et al.* 2007; CHIA *et al.* 2014). Transgenic animals expressing tagRFP::UtrCH in PVD neurons showed the reporter localized to defined subcellular compartments during development of the dendritic arbor. During the L2 larval stage tagRFP::UtrCH was primarily localized to the cell body, the 1°, 2°, and budding 3° branches (Figure 6A). During the subsequent larval stages, tagRFP::UtrCH remained strongly expressed in the cell body, but otherwise became successively more localized to distal (i.e. developing) branches, rather than proximal branches. For example, at the young adult stage, tagRFP::UtrCH was primarily localized to 4° branches, and less to 3° and 2° branches (Figure 6A). These results indicate that tagRFP::UtrCH, and by inference F-actin, localizes to extending branches. Conversely, an α-tubulin fusion (tagRFP::TBA-1) fusion that labels microtubules, specifically expressed in PVD neurons is primarily localized to axons and 1° dendrites of PVD neurons (Figure S6), as previously described (MANIAR *et al.* 2012). Thus, microtubules and F-actin occupy non-overlapping compartments in dendrites. Interestingly, this largely mutually exclusive localization of F-actin and microtubules is lost in mutants of the Menorin pathway. For example, mutations in *dma-1/LRR-TM, hpo-30/Claudin, sax-7/L1CAM*, or *tiam-1/GEF* result in a relocalization of the UtrCH reporter, and by inference F-actin, to more proximal dendrites (Figure 6B) with more dendritic endings lacking F-actin in *tiam-1/GEF* mutants (Figure 6C). Conversely, the tagRFP::TBA-1 fusion can be found in more distal dendrites in mutants of the Menorin pathway (Figure S6). These results suggest that the Menorin pathway establishes polarity and may function to localize F-actin to growing dendrites.

**Figure 6.**
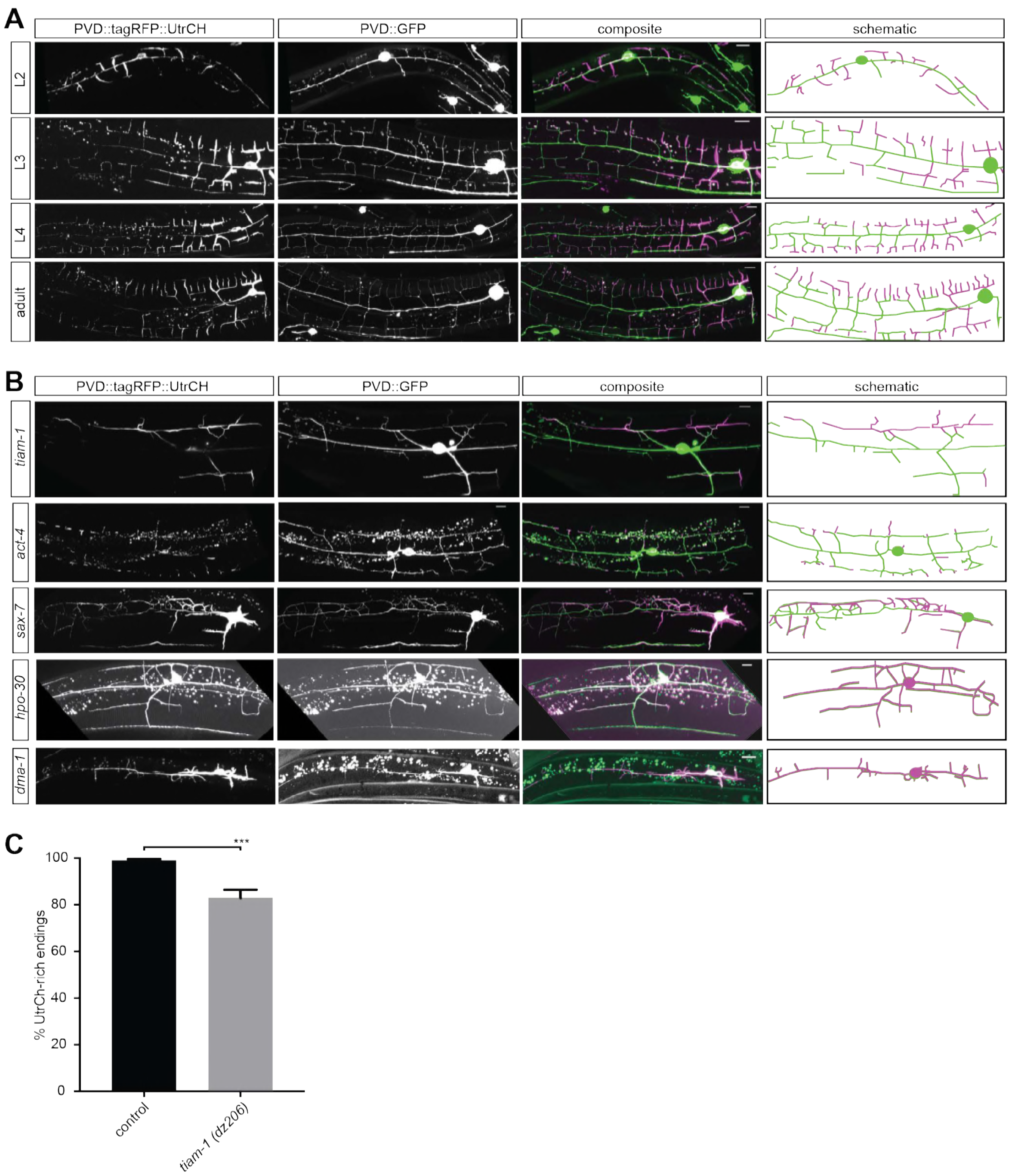
F-actin is localized to the leading edges of developing dendrites and requires the Menorin pathway for polarized localization. A. Fluorescent images of animals at different developmental stages carrying a F-actin reporter (tagRFP::UtrCH, left panels), a PVD cytoplasmic GFP reporter (second panels), merged images (third panel), and corresponding schematics (right panels). L2: second larval stage, L3: third larval stage, L4 fourth larval stage. Scale bars indicate 10μm. B. Fluorescent images of animals in different genetic backgrounds carrying a F-actin reporter (tagRFP::UtrCH, left panels), a PVD cytoplasmic GFP reporter (second panels), merged images (third panel), and corresponding schematics (right panels). Genotypes are indicated on the left. Scale bars indicate 10μm. C. Quantification of dendrite termini with F-actin rich staining in wild type control (n=10) and *tiam-1(tm1556)* mutant animals (n=13). Data are represented as mean ± SEM. Statistical comparisons were performed using the Mann-Whitney test. Statistical significance is indicated (***p < 0.005).

### TIAM-1/GEF forms a biochemical complex with the DMA-1/LRR-TM receptor and ACT-4/Actin

The remarkable similarity of phenotypes, namely a reduction of 4° branches, in the DMA-1(ΔPDZ) allele (Figure 1) and TIAM-1(GEF-only) rescued animals (Figure 5) suggested that DMA-1/LRR-TM and TIAM-1/GEF may interact directly through a PDZ interaction for the formation of 4° branches. We transfected DMA-1 (tagged with V5, DMA-1.V5) and TIAM-1 (tagged with HA, TIAM.HA) (Figure 7A) in human embryonic kidney cells (HEK293T) and conducted co-immunoprecipitation experiments from lysates of transiently transfected cells. We found that DMA-1.V5 efficiently co-immunoprecipitated TIAM-1.HA from lysates (Figure 7B). This interaction was dependent on the DMA-1 intracellular domain (ICD) and, specifically on the PDZ binding site, as the interaction was strongly reduced if either was removed (Figure 7C). Conversely, removing the PDZ domain from TIAM-1 (ΔPDZ) completely abrogated the interaction between DMA-1/LRR-TM and TIAM-1/GEF (Figure 7D). Additionally, removing the EVH1 domain from TIAM-1/GEF also compromised the interaction with DMA-1, albeit not to the same extent, suggesting that the EVH1 domain may serve some function, possibly as part of a larger complex. In conclusion, we propose that TIAM-1/GEF directly interacts through its PDZ domain with the PDZ binding site at the extreme C-terminus of DMA-1/LRR-TM, with a possible contribution of the EVH1 domain.

**Figure 7.**
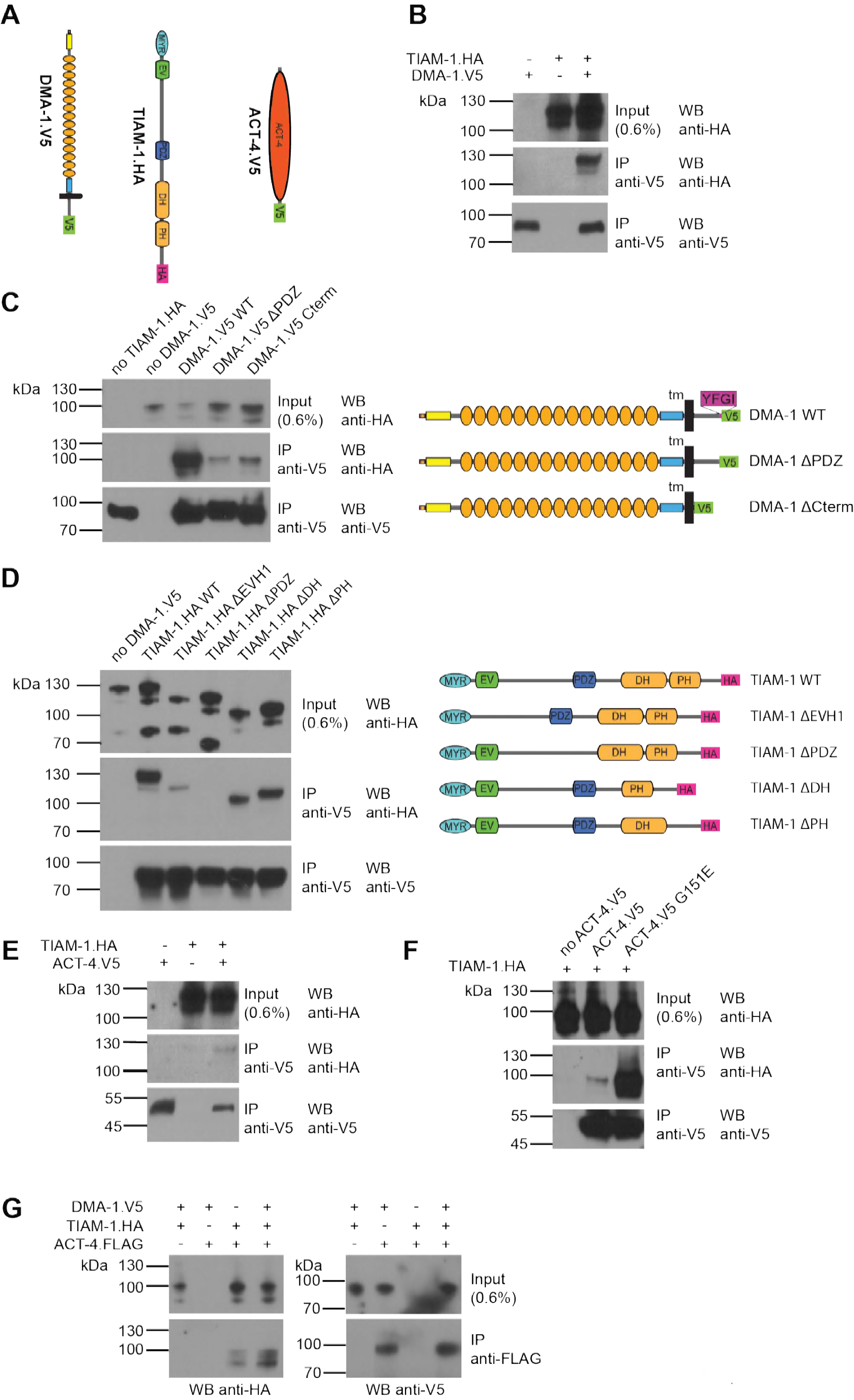
The DMA-1/LRR-TM, TIAM-1/GEF and ACT-4/Actin are part of the same biochemical complex. A. Schematic showing the topography of the DMA-1/LRR-TM single pass transmembrane receptor, the TIAM-1/GEF multidomain protein and the ACT-4/Actin. Schematics not to scale. Immuno tags (V5 and HA) used for co-immunoprecipitation experiments are shown. B. - F. Western Blots of co-immunoprecipitation experiments. Transfected constructs are indicated above the panels. Antibodies used for immunopreciptation (IP) and Western Blotting (WB) are indicated. A molecular marker is on the left. Note, that the two lo1wer panels are from a single Western blot, which was developed repeatedly with two different antibodies after stripping in all panels. Panel (B) investigates the interaction between TIAM-1 and DMA-1, and panels (C) and (D) are structure function analyses to delineate the domains required for the TIAM-1/DMA-1 interaction in TIAM-1 and DMA-1, respectively. Panels (E) and (F) analyze the interaction between TIAM-1 and wild type ACT-4 or mutant ACT-4(G151E), respectively.

Based on our genetic findings we next investigated the biochemical relationship of DMA-1/LRR-TM and TIAM-1/GEF with ACT-4/Actin to form a biochemical complex. We found that TIAM-1 co-immunoprecipitated ACT-4 from lysates (Figure 7E). Intriguingly, we found that the ACT-4 G151E missense mutation encoded by the *dz222* missense allele, which we isolated from our genetic screen, dramatically increased the strength of the biochemical interaction (Figure 7F). Deletion of individual domains TIAM-1 failed to abrogate the interaction with either wild type or G151E mutant ACT-4/Actin (Figure S6), suggesting that more than one domain in TIAM-1/GEF may act in a partially redundant fashion to bind ACT-4. Alternatively, a multiprotein complex could account for this observation. Surprisingly we found that ACT-4/Actin could also be precipitated with an antibody against DMA-1/LRR-TM, with the interaction possibly stronger in the presence of TIAM-1/GEF (Figure 7G). Collectively, we conclude that TIAM-1/GEF is part of a complex containing both the DMA-1/LRR-TM receptor and ACT-4/Actin, potentially directly linking the receptor to the cytoskeleton.

## Discussion

### DMA-1/LRR-TM regulates 3° and 4° branch formation through distinct mechanisms

Several lines of evidence suggest that signaling downstream of the DMA-1/LRR-TM receptor diverges into at least two molecularly distinct pathways with one functioning to establish 2° and 3° branches, whereas a pathway functioning at least partially in parallel serves to establish primarily 4° high order branches. First, removing the PDZ ligand motif of DMA-1/LRR-TM results in a decrease in 4° branches but not 2° or 3° branches. Conversely, a GEF-only construct of TIAM-1/GEF, which lacks the PDZ domain, rescues the 3° branching defects, but not 4° branching defects of *tiam-1* mutant animals. Thus, loss of either the PDZ domain in TIAM-1/GEF or PDZ ligand motif in DMA-1/LRR-TM results in loss of 4° branching. Furthermore, immunoprecipitation assays strongly indicate that TIAM-1/GEF and DMA-1/LRR-TM can interact directly through the PDZ motif at the C-terminus of DMA-1 and the PDZ-binding domain of TIAM-1. Since *tiam-1* - overexpression resulted in excessive branching, TIAM-1/GEF may promote branching in a dose-dependent manner. It can therefore be hypothesized that the PDZ interaction with DMA-1 recruits TIAM-1 to specific locations, where a locally higher concentration of TIAM-1 induces branching. Indeed, a recent study suggested that a TIAM-1::GFP reporter expressed in PVD is recruited to higher order dendritic branches by DMA-1/LRR-TM (ZOU *et al.* 2018). Collectively, these findings suggest that the PDZ-mediated interaction of TIAM-1/GEF and DMA-1/LRR-TM interaction is required for normal 4° branching.

An intriguing finding is that transgenic expression of a TIAM-1(GEF-only) construct lacking the PDZ interaction domain can rescue both the formation of 2° and 3° PVD branches. Conversely, DMA-1/LRR-TM lacking the cytoplasmic domain was still capable of rescuing 2° and 3° branching number (but not extension), suggesting that protein interactions of the DMA-1 transmembrane or the extracellular domain, possibly in conjunction with TIAM-1/GEF may mediate an alternative parallel downstream signaling path. Indeed, imaging experiments and biochemical assays suggest that (1) DMA-1/LRR-TM colocalizes with the HPO-30/Claudin and that (2) both proteins can form a biochemical complex through an interaction that is independent of the cytoplasmic domain of DMA-1. Moreover, genetic double mutant analyses show that *hpo-30* double mutants with *tiam-1* or *sax-7/L1CAM* display enhanced phenotypes consistent with functions in parallel pathways. How then could the *hpo-30*-dependent signaling pathway function mechanistically? A possible explanation comes from a recent study suggesting that HPO-30 can physically interact with WASP family verprolin-homologous protein (WVE-1/WAVE) regulatory complex (WRC) of actin regulators (ZOU *et al.* 2018), thereby controlling actin dynamics through this well-established mechanism. Surprisingly, neither loss of the actin regulator WSP-1/WASP nor the WASP family verprolin-homologous protein (WVE-1/WAVE) result in comparable dendrite patterning defects in *C. elegans* (LIAO *et al.* 2018) demonstrating that neither protein is individually required. Further experiments will be required to determine the precise mechanism by which DMA-1/LRR-TM, TIAM-1/GEF, and the WRC pattern PVD 2° and 3° dendrites.

### The DMA-1/LRR-TM may shape 4° dendrites by directly regulating F-actin

F-actin clearly plays an important role in the formation of PVD dendritic branches. First, depolymerizing F-actin *in vivo* results in severe defects in PVD patterning (HARTERINK *et al.* 2017). Second, F-actin is localized to the distal tips of developing PVD dendrites and appears mutually exclusive with microtubules. Third, we find that F-actin localization to distal dendrites depends on the presence of transmembrane receptors on PVD dendrites and their ligands. Lastly, RNAi-mediated gene knock down of other genes with possible or established roles as regulators of F-actin in developing axons such as *unc-44/Ankyrin* (OTSUKA *et al.* 1995), *unc-115/Ablim* (LUNDQUIST *et al.* 1998; STRUCKHOFF AND LUNDQUIST 2003), or *unc-70/β-spectrin* (HAMMARLUND *et al.* 2000), display similar defects in PVD dendrite patterning as *act-4/Actin* and *tiam-1/GEF* mutants (AGUIRRE-CHEN *et al.* 2011).

Our experiments show that the DMA-1/LRR-TM receptor is part of a biochemical complex with the TIAM-1/ GEF and ACT-4/Actin. How could this complex mediate 4° dendrite growth or patterning? One possibility is that the DMA-1/TIAM-1/ACT-4 complex is part of a molecular clutch mechanism, first proposed as a means to explain actin-dependent cellular movements by (MITCHISON AND KIRSCHNER 1988). These authors suggested that the force generated by polymerizing F-actin is transmitted through engagement of F-actin filaments with the membrane through transmembrane receptors. For example, transmembrane receptors of the Ig domain superfamily have been shown to couple extracellular interactions to cytoskeletal actin dynamics (reviewed in (GIANNONE *et al.* 2009)). In analogy, binding of the DMA-1/LRR-TM receptor by the extracellular L1CAM/MNR-1/LECT-2 ligand complex could result in recruitment of TIAM-1/GEF, ACT-4/Actin and DMA-1/LRR-TM in a complex, thereby directly linking the receptor to the cytoskeleton. The interaction between TIAM-1/GEF and ACT-4/Actin is likely specific, because *act-4(dz222)* mutant animals are viable without obvious phenotypes. This is stark contrast to the embryonic lethality observed upon continuous RNAi-mediated knockdown of *act-4* ((GOTTSCHALK *et al.* 2005), data not shown). Moreover, knock down of ACT-4/Actin by RNAi during larval stages on the one hand and, the G151E missense mutation in ACT-4/Actin encoded by the *act-4(dz222)* allele on the other hand result in similar phenotypes in PVD dendrites. In other words, loss of ACT-4 is as detrimental for dendrite patterning as a much stronger biochemical interaction between TIAM-1 and ACT-4(G151E). This suggests that the interaction between TIAM-1 and ACT-4, and possibly DMA-1/LRR-TM may have to be transient and, possibly, dynamic during dendrite patterning and, that the strong interaction between ACT-4(G151E) and TIAM-1/GEF sequesters actin from the polymerizing pool. These findings are reminiscent of a point mutation in β-spectrin, which by increasing the affinity of β-spectrin with actin one thousand fold, results in defects in drosophila dendrite arborization neuron (AVERY *et al.* 2017). Further experiments will be required to understand the dynamics of the TIAM-1/GEF-ACT-4/ACTIN interaction.

### *TIAM-1/*functions independently of its Rac1 GEF activity

Guanine nucleotide exchange factors such as TIAM-1 activate small GTPases by promoting the exchange of GDP for GTP. Tiam1 has been shown to serve important functions during nervous system patterning. For example, an enhancer trap line in mice shows severe malformations of the nervous system (YOO *et al.* 2012). Knock down and overexpression experiments further established functions for Tiam1 in axon growth cone formation and, activity-dependent remodeling of dendritic arbors *in vitro* and *in vivo* (KUNDA *et al.* 2001; TOLIAS *et al.* 2005; TOLIAS *et al.* 2007). Such functions may be conserved, since the drosophila homolog *still life/Tiam1* and *C. elegans tiam-1/GEF* function during synaptic and, axonal patterning downstream of the netrin receptor UNC-40/DCC (Deleted in Colorectal Cancer), respectively (SONE *et al.* 1997; DEMARCO *et al.* 2012). All known functions of Tiam1 appear dependent on the canonical enzymatic activity of exchanging GDP for GTP in small GTPases of the Rac1 type.

Unexpectedly, our findings imply that TIAM-1/GEF can function independently of its Rac1 GEF activity in shaping PVD dendrites. This conclusion is supported by three arguments. First, transgenic expression of a TIAM-1 (T548F) fully rescued the *tiam-1* mutant phenotype. Second, engineering the T548F point mutation into the *tiam-1* locus resulted in animals with PVD dendritic arbors that were indistinguishable from wild type animals. The T548F mutation is analogous to the S1216F mutation in UNC-73/Trio, which lacks Rac1 activity, but displays neuronal patterning defects (STEVEN *et al.* 1998). Moreover, the very mutation in *C. elegans* TIAM-1/GEF also shows no Rac1 GEF activity *in vitro* (DEMARCO *et al.* 2012). Lastly, single and double mutants of the small GTPases *mig-2/RhoG* and *ced-10/Rac1*, previously established as acting downstream of TIAM-1/GEF during axonal patterning in *C. elegans* (DEMARCO *et al.* 2012), displayed no defects PVD morphology. Similarly, a third Rac-like gene (*rac-2/3*) encoded in the *C. elegans* genome serves no function individually in PVD patterning. Taken together, these findings suggest that TIAM-1/GEF can function independently of Rac1 activity during PVD patterning.

To our knowledge, there is only one other known example of GEF-independent functions of a guanine nucleotide exchange factor. The DOCK 180 family member DOCK7 functions during nuclear migration by antagonizing TACC3, a protein known to coordinate microtubule polymerization (YANG *et al.* 2012). TACC3 is part of a protein complex conserved in *C. elegans*, consisting of *tac-1/TACC3, zyg-8/doublecortin*, and *zyg-9/XMAP215*, which is important for microtubule-dependent chromosome segregation during mitosis (BELLANGER AND GÖNCZY 2003; LE BOT *et al.* 2003; SRAYKO *et al.* 2003; BELLANGER *et al.* 2007). However, mutations in *tac-1/TACC3, zyg-8/doublecortin*, or *zyg-9/XMAP215* did not result in obvious defects in PVD patterning either in a wild type or *tiam-1/GEF* mutant background (data not shown). In conclusion, GEF-independent functions that directly modulate the cytoskeleton may be a more general and, previously underappreciated property of guanine nucleotide exchange factors.

## Material and Methods

### *C. elegans* strains and imaging

All strains were maintained using standard methods (BRENNER 1974). All experiments were performed at 20°C, and animals were scored as 1-day-old adults unless otherwise specified. The strains and mutant alleles used in this study are listed in the Supplementary Experimental Procedures. Fluorescent images were captured in live *C. elegans* using a Plan-Apochromat 40*/1.4 or 63x/1.4 objective on a Zeiss Axioimager Z1 Apotome. Worms were immobilized using 5mM levamisole and *Z* stacks were collected. Maximum intensity projections were used for further analysis.

For quantification of branching, synchronized starved L1 larvae were allowed to grow for 50 hrs (corresponding to late L4-young adulthood) at which time they were mounted. Fluorescent images of immobilized animals (1-5 mM levamisol, Sigma) were captured using a Zeiss Axioimager. *Z* stacks were collected and maximum projections were used for tracing of dendrites as described (SALZBERG *et al.* 2013). Briefly, all branches within 100 μm of the primary branch anterior to the cell body were traced, measured, and classified into primary, secondary, tertiary, quaternary, and ectopic tertiary branches using the NeuronJ plugin of the ImageJ 1.46r software. Statistical comparisons were conducted using one-sided ANOVA with the Tukey correction, the Kruskal-Wallis test, the Student t-test or the Mann-Whitney test as applicable using the (Prism [GraphPad]) software suite.

### Cloning of mutant alleles

Alleles of different genes were isolated during genetic screens for mutants with defects in PVD patterning. All screens were clonal F1 screens. Details on genetic screens, positional cloning and complementation tests are provided in Supplementary Experimental Procedures.

### Molecular biology and transgenesis

To establish DNA constructs used for rescue experiments or immunoprecipitation experiments, the respective cDNAs were cloned under control of either cell-specific promoters or promoters to drive expression in cell culture. Details for transgenesis, plasmid construction and a complete list of transgenic strains can be found in Supplementary Experimental Procedures.

### Author contributions

C. A.D.B, L.T., M.R., N.J.R.S., Y.S. and H.E.B. conceived, and C.A.D.B, L.T., M.R., N.R.S., and M.I.L.P. performed experiments. Y.S. was involved during inception of the project, isolation, cloning and initial characterization of several mutant alleles. C.A.D.B, L.T., M.R., and H.E.B. analyzed the data and prepared the manuscript with editorial input from all authors.

## Acknowledgments

We thank T. Boulin, S. Cook, A. Meléndez, R. Townley and members of the Bülow laboratory for comments on the manuscript and for discussion during the course of this work. We thank the Caenorhabditis Genome Center for strains (funded by NIH Office of Research Infrastructure Programs, P40OD010440); O. Hobert and K. Shen for reagents and discussions. This work was funded in part through the NIH (R21NS081505 to H.E.B; T32GM007288 and F31HD066967 to C.A.D.B.; F31NS100370 to M.R.; T32GM07491 to M.I.L.P.; P30HD071593 to Albert Einstein College of Medicine) and a Human Genome Pilot Project from Albert Einstein College of Medicine. N.J.R.S is the recipient of a Fulbright-Colciencias fellowship. H.E.B. is an Irma T. Hirschl/Monique Weill-Caullier research fellow.

## Supplementary Materials and Methods

### *Strain list*

#### Fluorescent reporter strains

PVD: *wdIs52 [PF49H12.4::GFP]* II (kind gift of David Miller)
PVD: *wyIs378* [Pser-2prom3::MYR-GFP + Pra6-3::MYR-mCherry] II (kind gift of Kang Shen)
PVD: *dzIs53* [PF49H12.4::mCherry]
Touch receptor neurons: *muIs32 [Pmec-7::GFP]* II

*Strains used in this study:*

NC1687: *wdIs52 II*
EB2981: *tiam-1*(*tm1556*) *I; wdIs52 II*
EB1278: *tiam-1*(*dz184*) *I; wdIs52 II*
EB2566: *tiam-1*(*dz206*) *I; wdIs52 II*
EB2982: *tiam-1*(*ok772*) *I; wdIs52 II*
EB2798: *wdIs52 II; hpo-30(ok2047) V*
EB1269: *wdIs52 II; hpo-30(dz189) him-5(e1490) V*
EB1279: *wdIs52 II; hpo-30(dz178) V*
EB2984: *wdIs52 II; act-4(dz222) X*
EB2985: *wdIs52 II; act-4(gk279371) X*
EB2986: *wdIs52 II; act-4(gk279385) X*
EB2987: *wdIs52 II; act-4(gk473333) X*
EB2988: *wdIs52 II; act-4(gk785720) X*
EB2730: *lect-2(rz2) wdIs52 II*
EB1564: *dma-1(tm5159)I; wdIs52 II*
EB1271: *wdIs52 II; mnr-1(dz175) V*
EB1899: *wdIs52 II; sax-7(nj48) IV*
EB1470: *kpc-1(gk8) I; wdIs52 II*
EB2989: *rab-10(ok1494) I; wdIs52 II*
EB1649: *wdIs52 II; dzIs43*
EB2990: *tiam-1(tm1556) I; wdIs52 II; dzIs43*
EB2991: *wdIs52 II; hpo-30(ok2047) V; dzIs43*
EB2992: *wdIs52 II; act-4(dz222) X; dzIs43*
CF702: *muIs32 II*
EB2993: *muIs32 II; act-4(dz222) X*
EB2994: *tiam-1(tm1556) I; muIs32 II*
EB2995: *muIs32 II; hpo-30(ok2047) V*
EB2996: *tiam-1(tm1556) I; wdIs52 II; mnr-1(dz175) V*
EB2997: *tiam-1(tm1556) I; wdIs52 II; sax-7(nj48) IV*
EB2998: *tiam-1(tm1556) dma-1(tm5159) I; wdIs52 II*
EB2999: *tiam-1(tm1556) kpc-1(gk8) I; wdIs52 II*
EB3000: *tiam-1(tm1556) rab-10(ok1494) I; wdIs52 II*
EB3001: *tiam-1(tm1556) I; lect-2(rz2) wdIs52 II*
EB3002: *tiam-1(tm1556) I; wdIs52 II; hpo-30(ok2047) V*
EB3003: *tiam-1(tm1556) I; wdIs52 II; act-4(dz222) X*
EB3004: *wdIs52 II; sax-7(nj48) IV; hpo-30(ok2047) V*
EB3005: *dma-1(tm5159) I; wdIs52 II; hpo-30(ok2047) V*
EB2808: *dzIs53 ddIs290 II; him-5(ok1896) V*
EB3006: *dzIs53 ddIs290 II; hpo-30(ok2047) V*
EB3007: *dzIs53 ddIs290 II; act-4(dz222) X*
EB3171 : *tiam-1(tm1556) I; dzIs53 ddIs290 II*
EB2724: *lect-2(dz249)[lect-2::mNG^^^3xFlag]) II*
EB3008: *lect-2(dz249)[lect-2::mNG^^^3xFlag]) II; hpo-30(ok2047) V*
EB3009: *lect-2(dz249)[lect-2::mNG^^^3xFlag]) II; act-4(dz222) X*
EB3172: *tiam-1(tm1556) I; lect-2(dz249)[lect-2::mNG*3xFlag]) II*
EB3010: *wyEx4286; tiam-1(tm1556) I*
EB3011: *wyEx4286; hpo-30(ok2047) V*
EB3012: *wyEx4286; act-4(dz222) X*
EB3013: *tiam-1(dz264)[TIAM-1 T548F] I; wdIs52 II*
EB3014: *tiam-1(dz265)[TIAM-1 T548F] I; wdIs52 II*
EB2960: *dzEx1566; wdIs52 II*
EB2980: *dzEx1566; tiam-1(tm1556) I; wdIs52 II*
EB3017: *dzEx1566; wdIs52 II; act-4(dz222) X*
EB3058: *dzEx1566; dma-1(tm5159) I; wdIs52 II*
EB3167: *dzEx1566; wdIs52 II; hpo-30(ok2047) V*
EB3175: *dzEx1566; wdIs52 II; sax-7(nj48) IV*
EB2975: *dzEx1569; wdIs52 II*
EB3062: *dzEx1569; dma-1(tm5159) I; wdIs52 II*
EB3063: *dzEx1569; tiam-1(tm1556) I; wdIs52 II*
EB3168: *dzEx1569; wdIs52 II; hpo-30(ok2047) V*
EB3169: *dzEx1569; wdIs52 II; act-4(dz222) X*
EB3170: *dzEx1569; wdIs52 II; sax-7(nj48) IV; him-5(ok1869) V*
EB2871: *dzIs53 II; act-4(dz222) X*
EB2933: *dzEx1545; dzIs53 II; act-4(dz222) X*
EB2936: *dzEx1548; dzIs53 II; act-4(dz222) X*
EB2940: *dzEx1552; dzIs53 II; act-4(dz222) X*
EB2953: *tiam-1(dz206) I; wdIs52 II; act-4(dz222) X*
EB2693: *dma-1(dz244[dma-1::mNG^^^3xFlag]) I; dzIs53 II*
EB2945: *dzEx1555; wdIs52 II*
EB3015: *dzIs95*
EB3016: *dzIs95; wdIs52 II; act-4(dz222) X*
EB3176: *dma-1(tm5159) I; wdIs52 II; dzEx1571*
EB3183: *dma-1(dz266[Y600Amber]) I; wdIs52 II*

### *Transgenic strains*

A complete list of all transgenic strains created for this study is shown below. Additional details are provided in the lists of transgenic strains and plasmids.

#### PVD specific rescue of *dma-1* PVD dendrite branching defect

Complete rescue of the *dma-1* mutant phenotype was achieved in transgenic animals carrying a plasmid where the *dma-1* cDNA was driven under control of a shorter version of the previously reported *ser2prom3* promoter (TSALIK et al 2003). (The new promoter version *ser2prom3s)* encompasses 1825bp upstream of the predicted promoter for the C02D4.2d isoform.

For the deletion of the intracellular domain in *dma-1* (ΔICD) a mCherry fluorescent tag was inserted in frame after E532 leaving a protein devoid of the last 71 residues (see plasmid list below). Both constructs were injected individually at 5 ng/μl into *dma-1(tm5159) I; wdIs52 II*, together with the *Pmyo-3::tagRFP* marker (at 50 ng/μl) for the injection mix containing the entire *dma-1* cDNA or *Pmyo-2::mCherry* (at 50 ng/μl) for the *dma-1* (ΔICD) version. The final injection mix was brought to 100 ng/μl of DNA concentration with *pBluescript*.

#### Heterologous rescue of *tiam-1* PVD dendrite branching defect

The *tiam-1* cDNA was cloned under control of heterologous promoters: hypodermal *Pdpy-7* (Gilleard et al., 1997), body wall muscle *Pmyo-3* (Okkema et al., 1993), pan-neuronal *Prab-3* (NONET et al., 1997), and a PVD-specific *Pser-2prom3* promoter (Altun-Gultekin et al., 2001). All constructs were injected at 5 ng/μl into *tiam-1(tm1556) I; wdIs52 II*, together with the *Pmyo-3::mCherry* marker at 50 ng/μl and *pBluescript* at 50 ng/μl.

#### Heterologous rescue of *act-4* PVD dendrite branching defect

The *act-4* (or *act-1)* genomic DNA was cloned under control of heterologous promoters: hypodermal body wall muscle *Pmyo-3* (Okkema et al., 1993) and PVD-specific *Pser-2prom3* (Altun-Gultekin et al., 2001). All constructs were injected at 5 ng/μl into *wdIs52 II; act-4(dz222) X*, together with the *Pelt-2::gfp* marker at 5 ng/μl and *pBluescript* at 90 ng/μl.

##### Transcriptional reporter of *act-4*

The *act-4* transcriptional reporter promoter was constructed by cloning 2.6kb upstream of the predicted *act-4* translational start site (based on Figure S1) into *pPD95.75-NLS::mCherry*, and injected at 5 ng/μl into *wdIs52 II* animals, together with the *Pmyo-2::mNG* marker at 5 ng/μl and *pBluescript* at 90 ng/μl.

##### Translational reporter of *act-4*

The *act-4* cDNA was tagged at the C-terminus with tagRFP and expressed under the *Pser-2prom3* promoter (Okkema et al., 1993 TSALIK et al., 2003). It was injected at 5 ng/μl into *wdIs52 II; act-4(dz222)* animals together with the *Pmyo-2::mNG* marker at 10 ng/μl and *pBluescript* at 85 ng/μl.

##### tagRFP::UtrCH and *tagRFP::tba-1* reporters

UtrCH and *tba-1* were tagged with tagRFP and expressed under *Pser-2prom3* promoter. All constructs were injected at 5 ng/μl into *wdIs52 II*, together with the *Pmyo-3::tagBFP* marker at 10 ng/μl and *pBluescript* at 85 ng/μl

### *Generation of genome-edited strains through CRISPR-Cas9*

For introducing mutations onto endogenous loci of *dma-1(Y600Amber)* and *tiam-1(T548F)*, the CRISPR co-conversion as described in Arribere et al 2014 was utilized. Briefly, 2 appropriate sgRNA sequences were identified for each gene and introduced into Cas9-sgRNA expression vector pDD162, while repair oligos were ordered from IDT. Both Cas9-sgRNA constructs and repair oligo were injected into N2 animals along with Cas9-sgRNA and repair oligo for *dpy-10(cn64)* conversion. Rollers *(dpy-10(cn64)/+ animals)* from the F1 generation were selected and genotyped using restriction digest engineered on the repair oligo. Wildtype F2 animals were then selected from positive F1 animals to segregate the intended edit from *dpy-10(cn64)*.

For C-terminal mNeon Green knock-in of *dma-1*, we followed the protocol for self-excising electing cassette as described in Dickinson et al 2015. Briefly, 700bp sequences flanking the C-terminal of *dma-1* were amplified from genomic DNA and inserted into pDD268 (pdma-1repair). NC1687 worms were then injected with pdma-1repair and Cas9-sgRNA for dma-1 along with pmyo-2::mCherry, prab-3::mCherry and pmyo-3::mCherry. F1 roller worms that survived hygromycin treatment and display no mCherry fluorescent were selected. Upon confirmation of mNG knock-in visually, L1 worms are subjected to 34C heat shock for 4 hours to remove selection markers, which is presented as non-rolling F1s.

#### sgRNA sequences

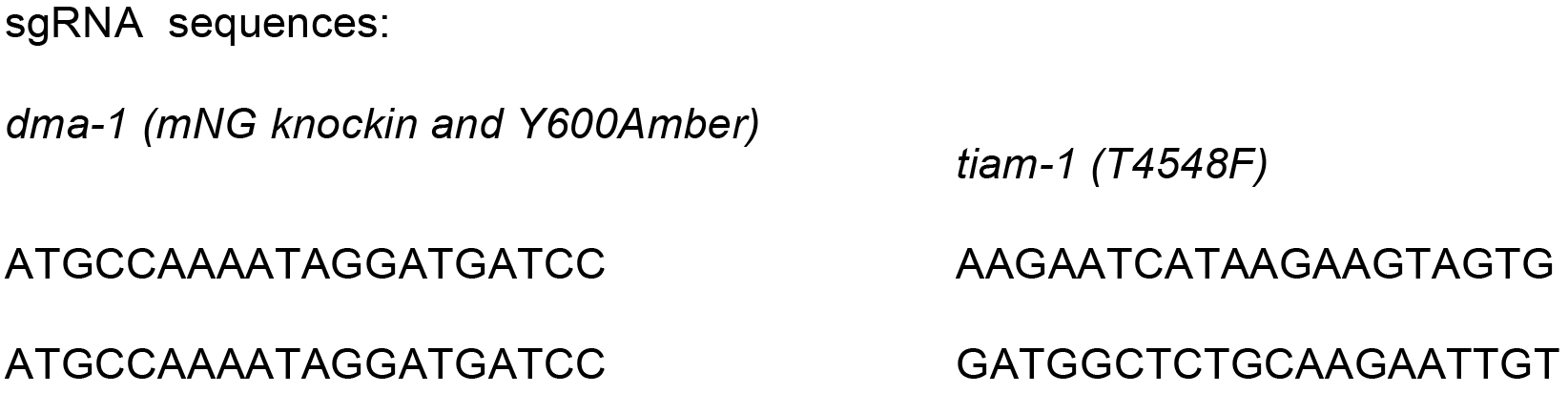
Problem-solving strategies that separate qualitative/pictorial steps from mathematical steps.

#### Repair oligo sequences

##### dma-1 (Y600Amber)

TGCAAATAATCACACGGATTTAGATTTTTGAAACATTTGAAACTCTCAAAAGGATATTTATTATCTAGACTAAGAGGAACCTGGCTTCGGAGGTGCTGGTGGAATCAATGGAGGAGTGGCTTTAAATGTTTCAGTACTAACCAAGAATGG

##### tiam-1 (T4548F)

TGTCAATAAGAAAACGGGCTGTGCAAAGTTGGCGATGGCTCTGCAAGAATTGTTAGTCTTTGAGAAGAAATATGTCAGCGATCTTCGAGAGGTAAGAGATCCCAAAAAGTTATAGAATTGAATAATTTACGATTTCAGATGA

### *Details of genetic screens and positional cloning*

In several F1 clonal genetic screens for mutants with defects in PVD dendrite arborization, we isolated one allele of *dma-1* (*dz181*), two alleles of *hpo-30* (*dz178* and *dz189*), and two alleles of *tiam-1* (*dz184* and *dz206*). One allele of *act-4* (*dz222*) was isolated in an enhancer screen of the hypomorphic *kpc-1(gk333538)* allele. In addition, we obtained putative null deletion alleles for *dma-1* (*tm5156*), *hpo-30* (*ok2047*), and *tiam-1* (*tm1556* and *ok772*). Details for cloning of individual point mutations is provided below.

***tiam-1(dz184):*** A SNP-mapping-WGS approach (DOITSIDOU et al., 2010) was used to map *tiam-1(dz184)* between 3Mb and 6Mb of chromosome I. Using SNP mapping approaches (DAVIS et al., 2005), the mutation further mapped to a physical interval between 4Mb and 5.48Mb. This region contained only one nonsense mutation. The mutation was confirmed by Sanger sequencing of the original isolate, confirming a non-sense C to T mutation in the *tiam-1(dz184)* allele. Additionally, the *tiam-1(dz184)* mutant showed non-complementation with all other loss-of-function alleles and its phenotype was not different from the *dz206, tm1556 and ok772* deletion alleles (see List of complementation tests below). Therefore, our genetic and molecular evidence suggests that all alleles used in this study are likely complete loss of function alleles of *tiam-1.*

***tiam-1(dz206):*** Complementation tests with a deletion allele of *tiam-1* showed noncomplementation between *tiam-1(tm1556)* and *tiam-1(dz206)*. The mutation was further confirmed by Sanger sequencing of the original isolate, identifying a non-sense mutation C to T mutation in *dz206* in the *tiam-1* locus (see List of *tiam-1* mutant alleles below).

*hpo-30(dz178):* A SNP-mapping-WGS approach (DOITSIDOU et al., 2010) was used to map *hpo-30(dz178)* between 3Mb and 5Mb of chromosome V. This region contained only 1 candidate mutation. The mutation was confirmed by Sanger sequencing of the original isolate, confirming a missense C to T mutation in *hpo-30(dz178)*. Additionally, the *hpo-30(dz178)* phenotype was not different from the *dz178 and ok2047* deletion allele. Therefore, our genetic and molecular evidence suggests that all alleles used in this study are likely complete loss of function alleles of *hpo-30*.

***hpo-30(dz189):*** Complementation tests with the other *loss-of-function* allele of *hpo-30* showed non-complementation between *hpo-30(dz178)* and *hpo-30(dz189)*. The mutation was further confirmed by Sanger sequencing of the original isolate, identifying a non-sense mutation C to T mutation in *hpo-30(dz189)* (see List of *hpo-30* mutant alleles below).

*act-4(dz222):* The *dz222* allele was identified in a genetic screen as a recessive enhancer of the *kpc-1(gk333538)* hypomorphic allele. A SNP-mapping-WGS approach (DOITSIDOU et al., 2010) was used to map *act-4(dz222)* between 4Mb and 6Mb of chromosome X. This region contained 5 candidate mutations, including non-sense, missense, frameshift and splice site mutations. The mutant defect in *dz222* was phenocopied by RNAi-mediated postembryonic gene knock down of *act-4* and, rescued by transgenic expression of *act-4* under control of heterologous promoters. Sequencing of the original isolate identified a G to A missense mutation in *act-4(dz222)*, resulting in glycine to glutamic acid change at position 151 in ACT-4. Finally, when *act-4(dz222)* allele was placed over a deficiency the phenotype was not enhanced. Therefore, our genetic and molecular evidence suggests that the *act-4(dz222)* allele used in this study is likely complete loss of function allele of *act-4* in the context of PVD dendrite morphogenesis.

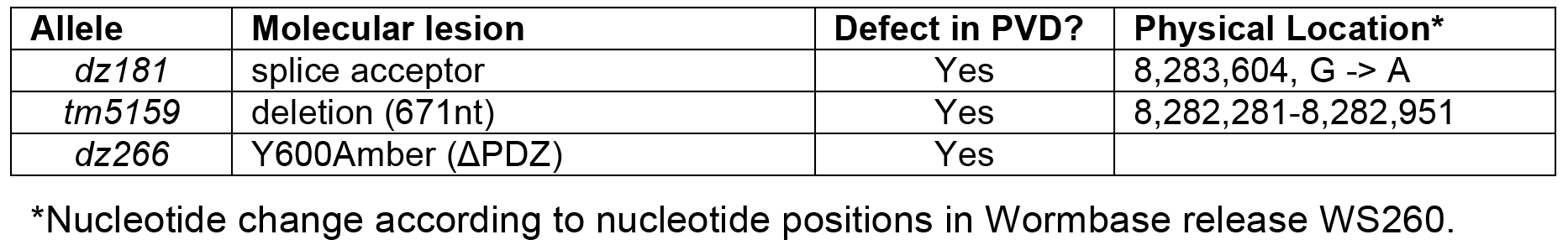
List of mutant *dma-1* alleles

List of mutant *hpo-30* alleles

List of mutant *hpo-30* alleles
| Allele | Molecular lesion | Defect in PVD? | Physical Location* |
| --- | --- | --- | --- |
| *dz178* | S155F | Yes | 12,309,368 |
| *dz189* | Q168X | Yes | 12,309,330 |
| *ok2047* | Deletion (1,294bp) | Yes | 12,309,724-12,311,017 |

\*Nucleotide change according to nucleotide positions in Wormbase release WS252.

List of mutant *tiam-1* alleles

List of mutant *tiam-1* alleles
| Allele | Molecular lesion | Defect in PVD? | Physical Location* |
| --- | --- | --- | --- |
| *dz184* | Q303X | Yes | 4,422,065 |
| *dz206* | Q543X | Yes | 4,424,113 |
| *dz264* | T548F | No | 4,424,128-4,424,129 |
| *dz265* | T548F | No | 4,424,128-4,424,129 |
| *tm1556* | Deletion (851bp) | Yes | 4,423,487-4,424,337 |
| *ok772* | Deletion (838bp) and insertion (18bp) | Yes | 4,424,394-4,425,231 |

\*Nucleotide change according to nucleotide positions in Wormbase release WS252.

List of mutant *act-4* alleles

List of mutant *act-4* alleles
| Allele | Molecular lesion | Defect in PVD? | Physical Location* |
| --- | --- | --- | --- |
| *dz222* | G151E (M03F4.2a) | Yes | 4,963,194 |
| *gk279371* | L9H (M03F4.2a) | No | 4,960,687 |
| *gk279385* | G246R (M03F4.2a) | No | 4,963,478 |
| *gk473333* | A30T (M03F4.2a) | No | 4,962,830 |
| *gk785720* | E4K (M03F4.2c) | No | 4,962,048 |

\*Nucleotide change according to nucleotide positions in Wormbase release WS252.

List of Complementation tests for *tiam-1*

List of Complementation tests for *tiam-1*
| genotypes | % of defective PVDs | N | Results |
| --- | --- | --- | --- |
| +/+ (*wdIs52* (control)) | 0 | 100 | Complementation |
| *dz184/tm1556* | 100 | 100 | Non-complementation |
| *dz184/ok772* | 100 | 100 | Non-complementation |
| *dz184/+* | 0 | 100 | Complementation |
| *dz206/tm1556* | 100 | 40 | Non-complementation |
| *dz206/+* | 0 | 40 | Complementation |

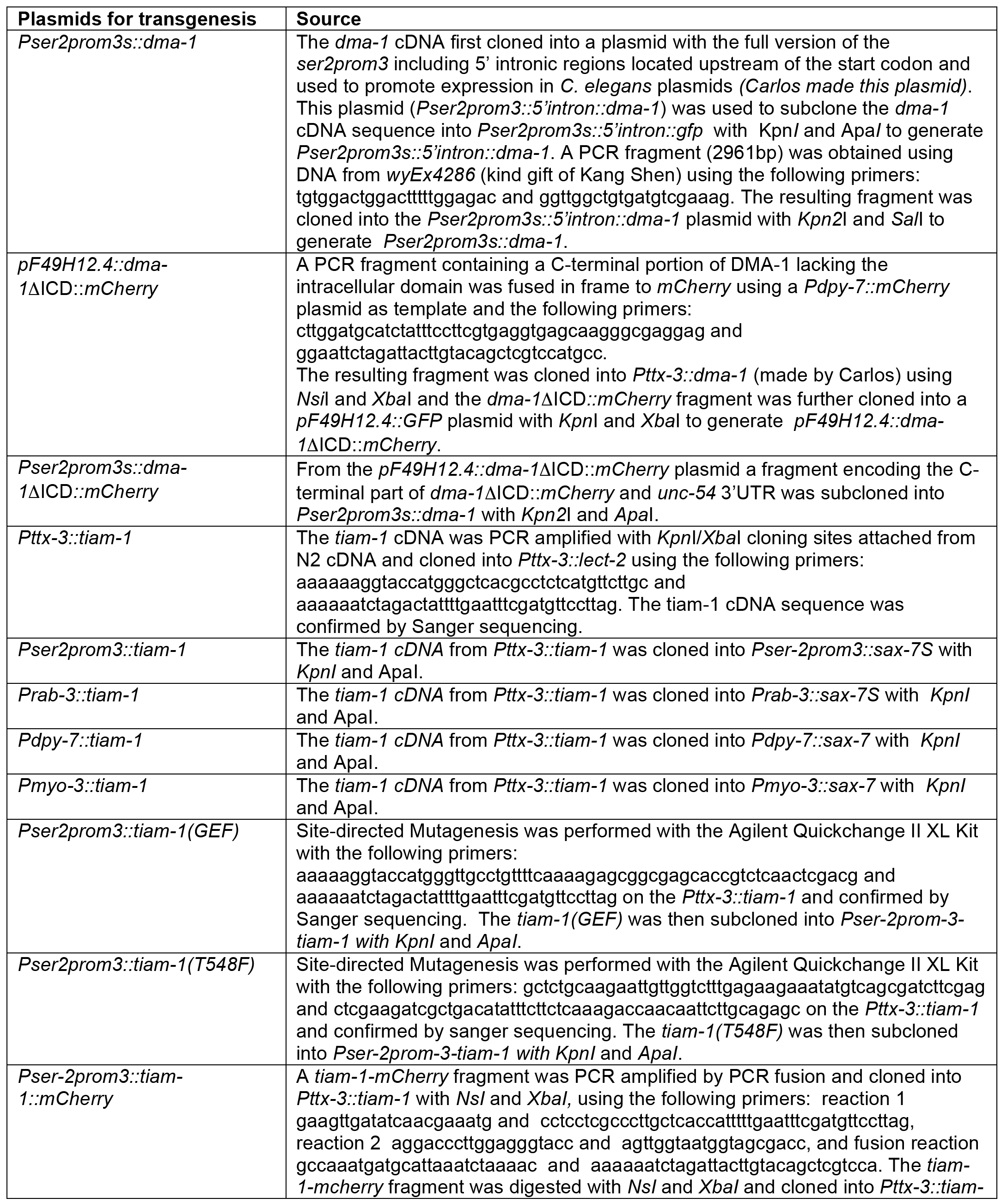
List of plasmids used in transgenic experiments

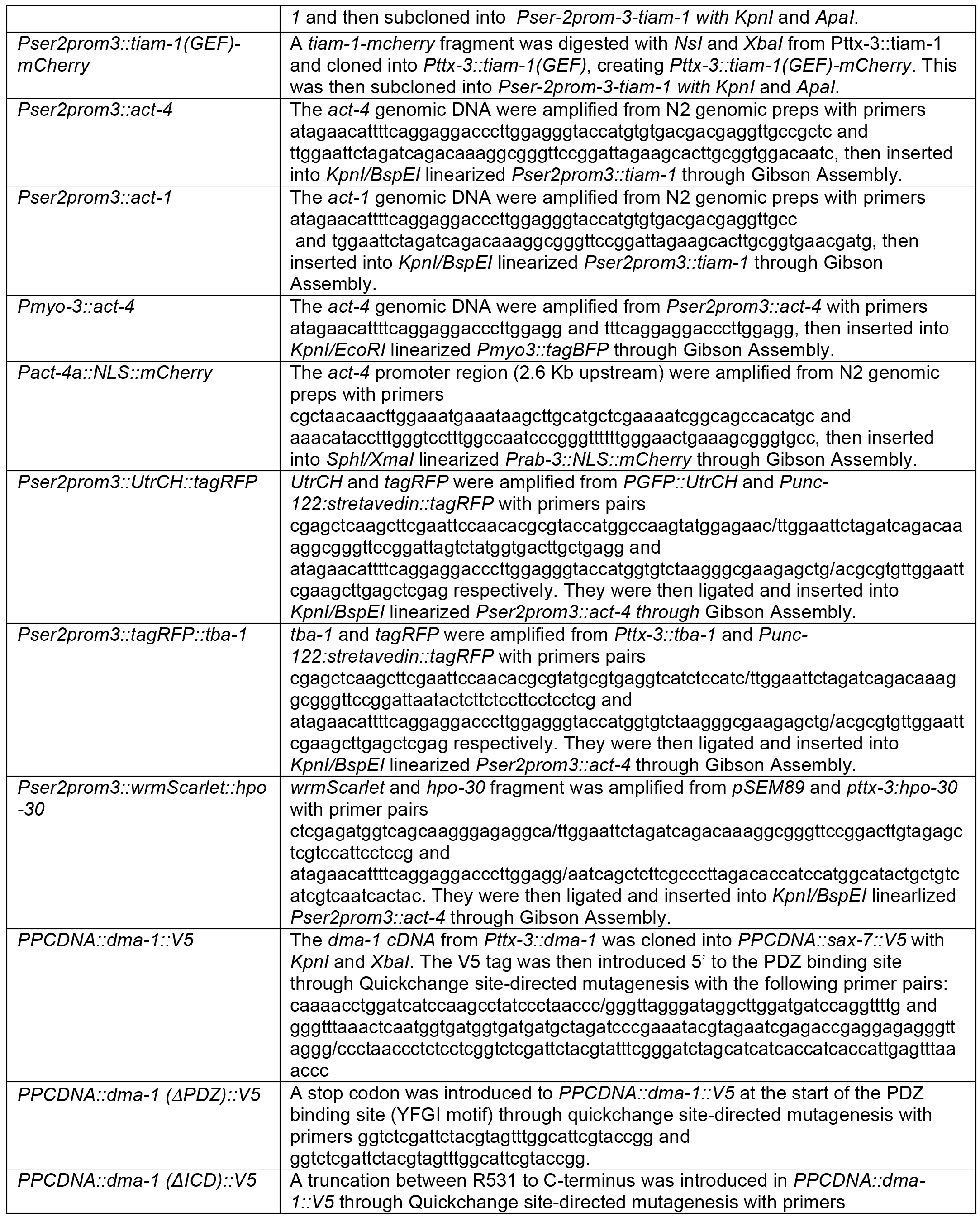

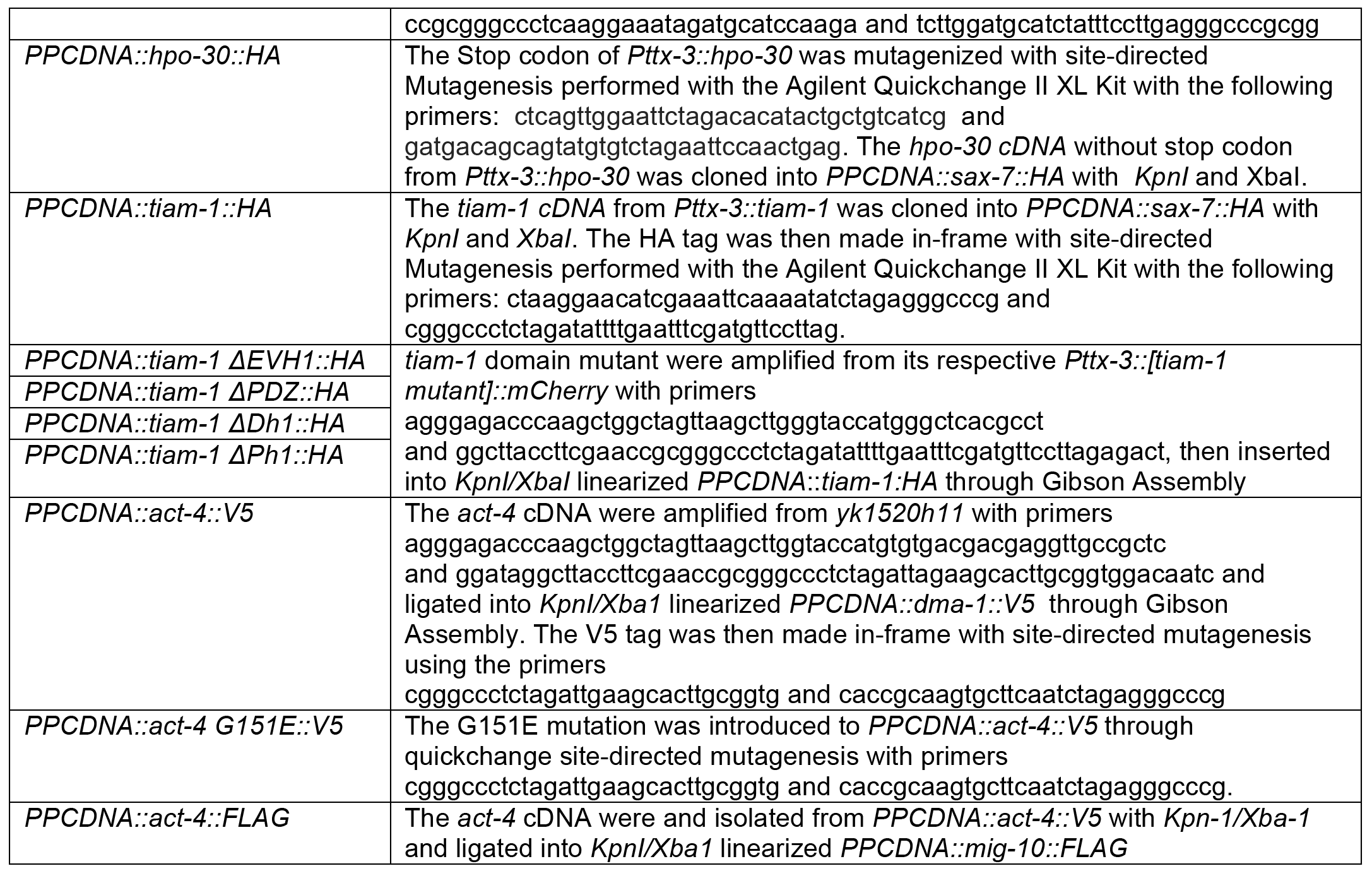

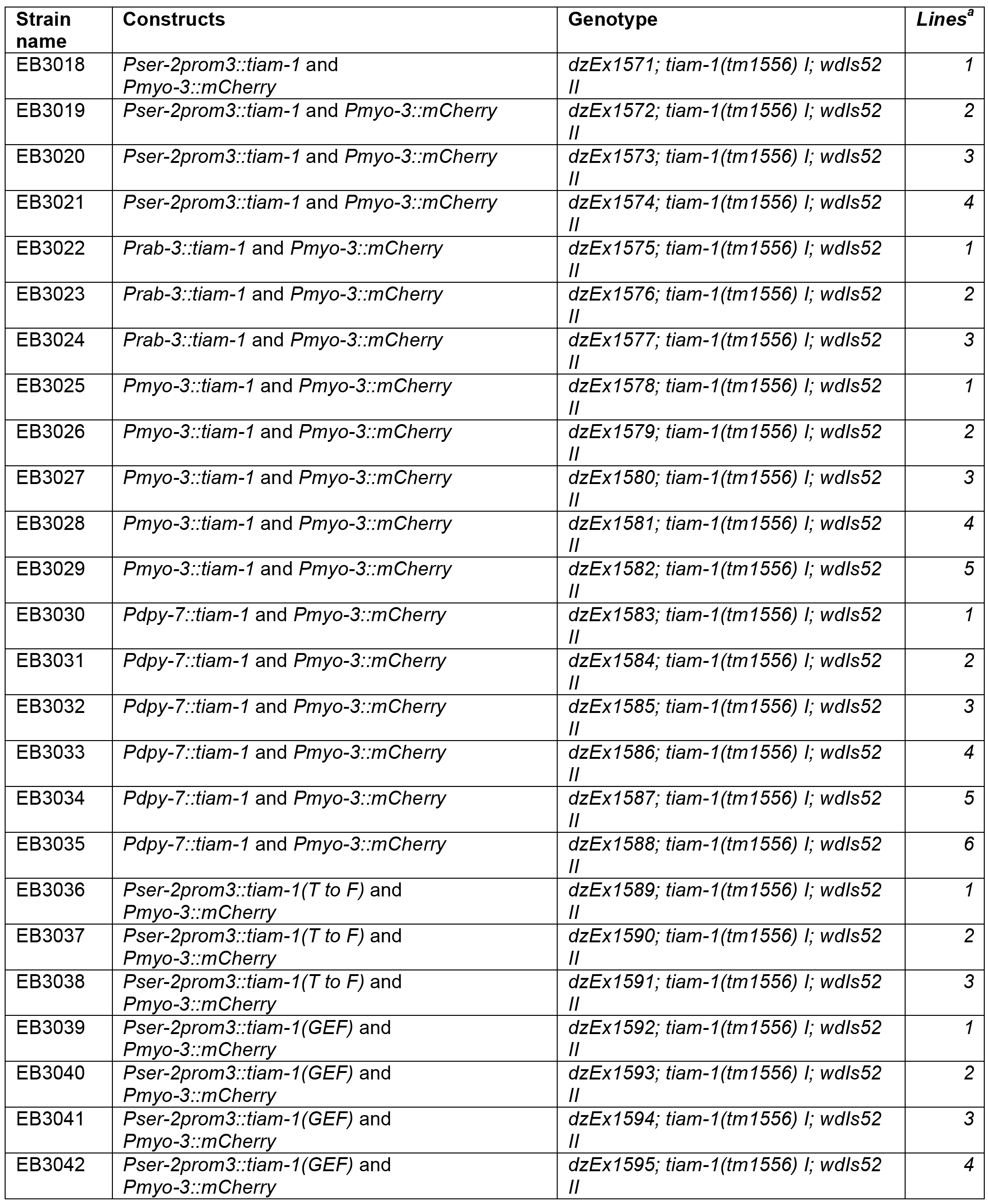
List of transgenic strains with genotypes and strain names

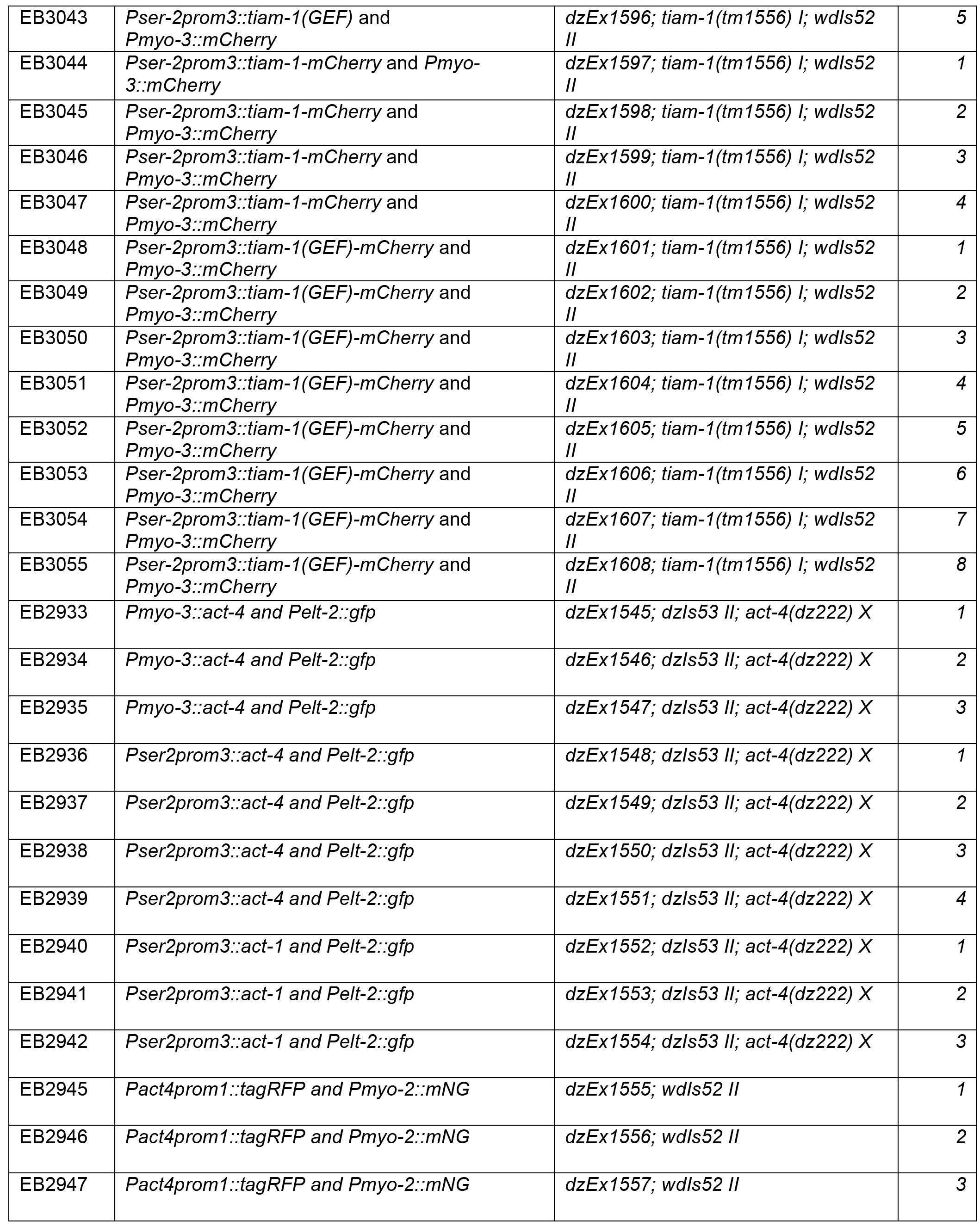

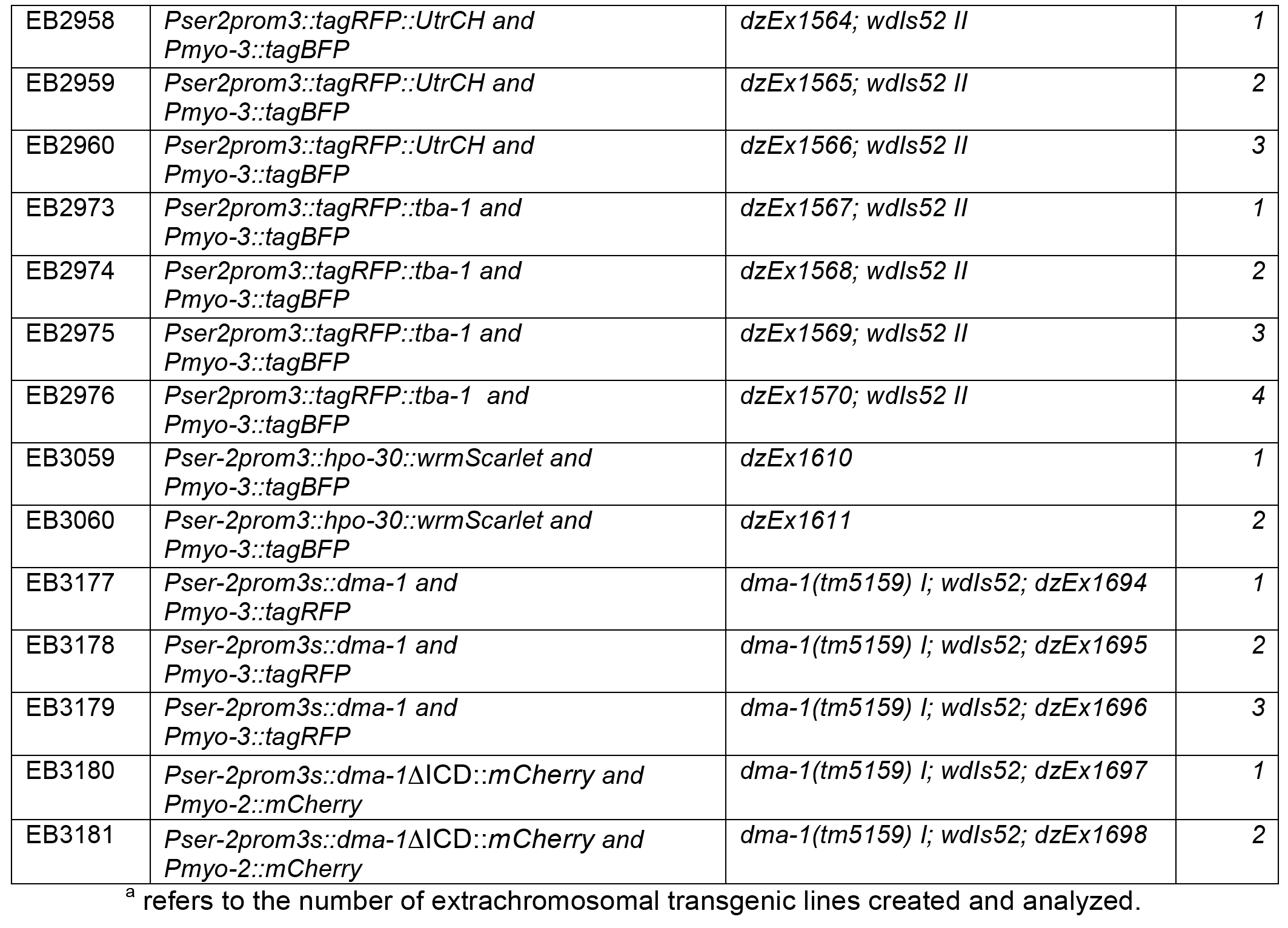

## Supplementary Figures

**Figure S1.**
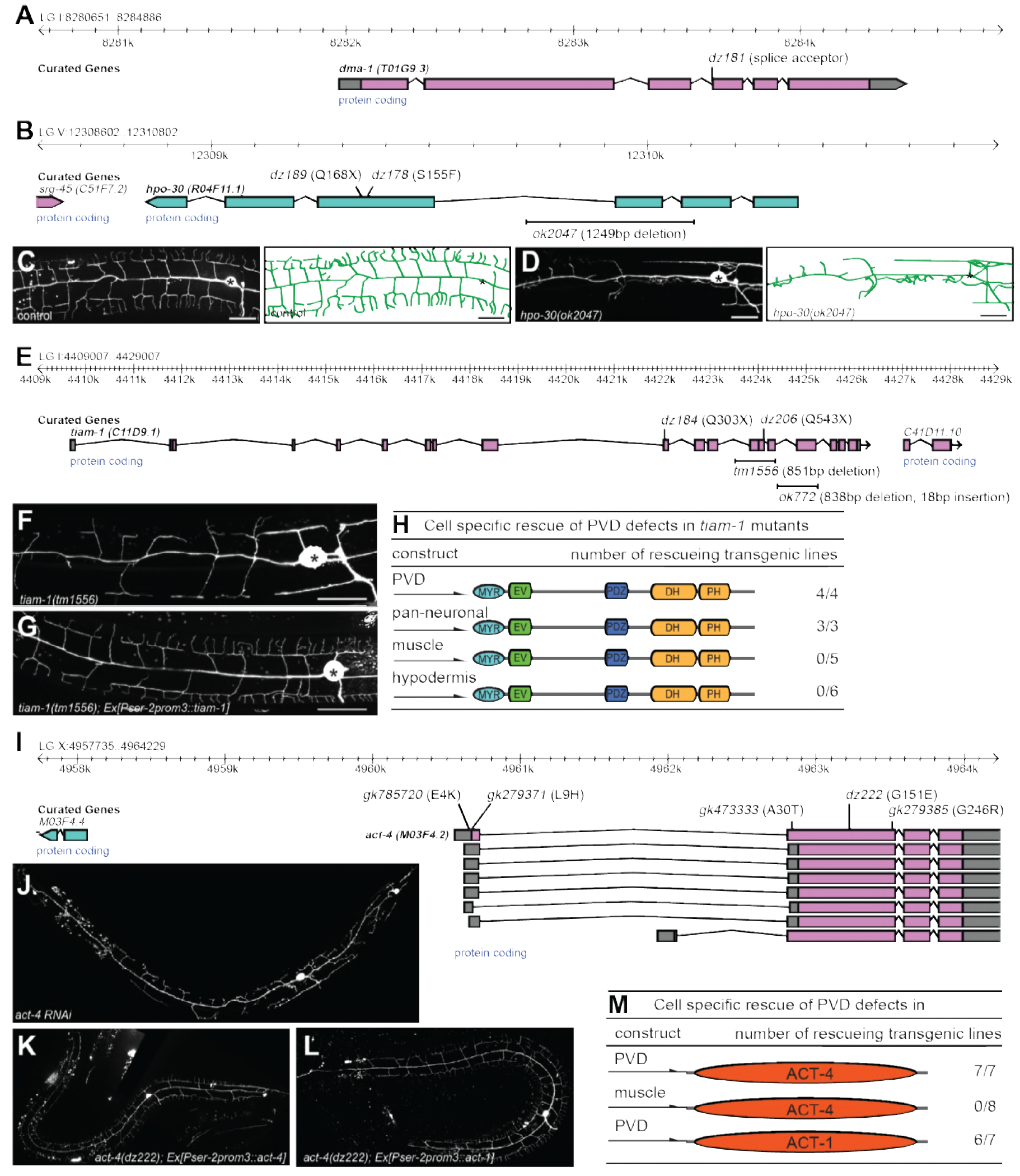
Genes functioning cell-autonomously in PVD somatosensory neurons. A. - B. Genomic environs of the indicated genes with the physical location on the respective linkage groups (LGs) are shown. The exon-intron structure is indicated, as is the direction of transcription. Alleles and the resulting molecular changes are shown above (for point mutants) and below (for deletions) the gene structure, respectively. C. - D. Fluorescent images of PVD (left panels) and schematics (right panels) of wild type control (C) and *hpo-30(ok2047)* mutant animals (D). PVD is visualized by the *wdIs52 [F49H12.4::GFP]* transgene and, anterior is to the left and dorsal is up in all panels; scale bars indicate 20μm. E. Genomic environs of *tiam-1* with the physical location on linkage group I are shown. The exon-intron structure is indicated as well as the direction of transcription. Alleles and the resulting molecular changes are shown above (for point mutants) and below (for deletions) the gene structure, respectively. F. - G Fluorescent images of PVD of *tiam-1(tm1556)* animals without (F) and with a transgene expressing a wild type TIAM-1 cDNA under control of the PVD specific *Pser-2prom3* promoter (TSALIK et al., 2003) (G). H. Summary of transgenic rescue experiments of *tiam-1(tm1556)* mutant animals. Cell-specific promoters are shown on the left (PVD: *Pser-2prom3* (TSALIK et al., 2003); pan-neuronal: *Prab-3* (NONET et al., 1997); *Pmyo-3* muscle: (OKKEMA et al., 1993); and *Pdpy-7* hypodermis (GILLEARD et al., 1997)). The number of independent transgenic lines out of the total number of transgenic lines obtained is shown on the right. I. Genomic environs of *act-4* with the physical location on linkage group X are shown. The exon-intron structure is indicated as well as the direction of transcription. Alleles and the resulting molecular changes are shown above the gene structure. J. - L. Fluorescent images of PVD in wild type animals after RNAi mediated gene knock down of *act-4* and, of *act-4(dz222)* mutant animals carrying a transgene expressing a wild type ACT-4 (K) or ACT-1 (L) genomic DNA, respectively, under control of the PVD specific *Pser-2prom3* promoter (TSALIK et al., 2003). M. Summary of transgenic rescue experiments of *act-4(dz222)* mutant animals with ACT-4 or ACT-1, respectively. Cell-specific promoters are shown on the left (PVD: *Pser-2prom3* (TSALIK et al., 2003); *Pmyo-3* muscle: (OKKEMA et al., 1993)). The number of independent transgenic lines out of the total number of transgenic lines obtained is shown on the right.

**Figure S2.**
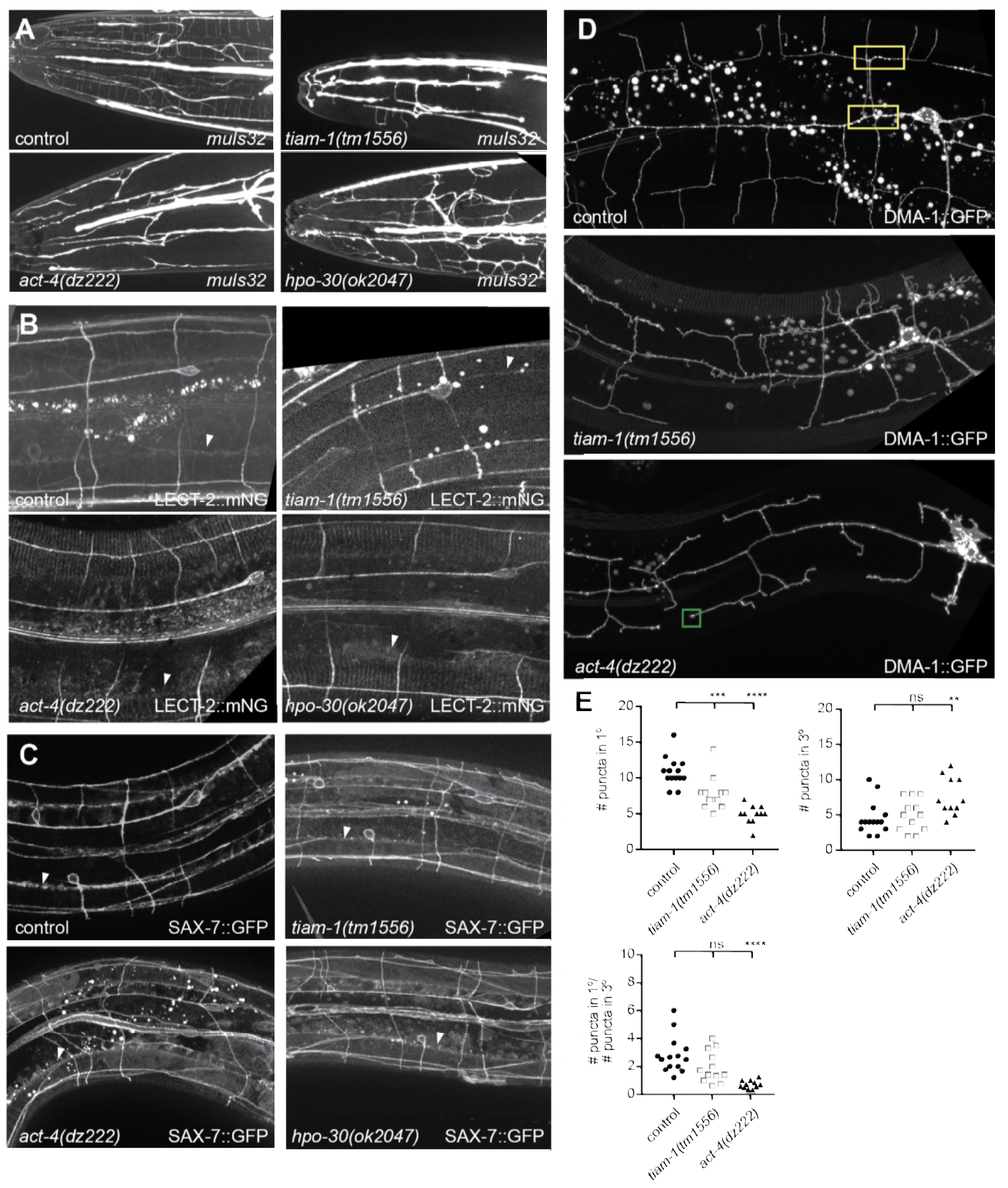
Effects of *hpo-30/Claudin, tiam-1/GEF* and *act-4/Actin* on localization of LECT-2:mNG, SAX-7::GFP and DMA-1::GFP. A. Fluorescent images of animals expressing GFP in FLP neurons *(muIs32 Is[Pmec-7::GFP])* in the indicated genotypes. Anterior is to the left and scale bars indicate 20μm. B. - D. Fluorescent images of animals expressing a LECT-2::mNeonGreen fusion at endogenous levels (a functional knock in *(lect-2(dz249)* [lect-2::mNG^^^3xFlag](DIAZ-BALZAC et al., 2016))(B), a functional fosmid-based SAX-7::GFP reporter (C), and a functional DMA-1::GFP reporter (D)(LIU AND SHEN, 2011). Arrowheads indicate the localization of LECT-2::mNG or SAX-7::GFP in a stripe at edge of the lateral hypodermis, where hypodermis and muscle abut. The control image (B) is identical to Figure 4B and shown for comparison only. E. Quantification of DMA-1::GFP puncta in the genotypes indicated. n=14 for control, n=12 for *tiam-1(tm1556)*, and n= 11 for *act-4(dz222)*. Data are represented as mean ± SEM. Statistical comparisons were performed using one-sided ANOVA with the Tukey correction. Statistical significance is indicated (ns: not significant, **p<0.05, ***p<0.005, ****p < 0.0005).

**Figure S3.**
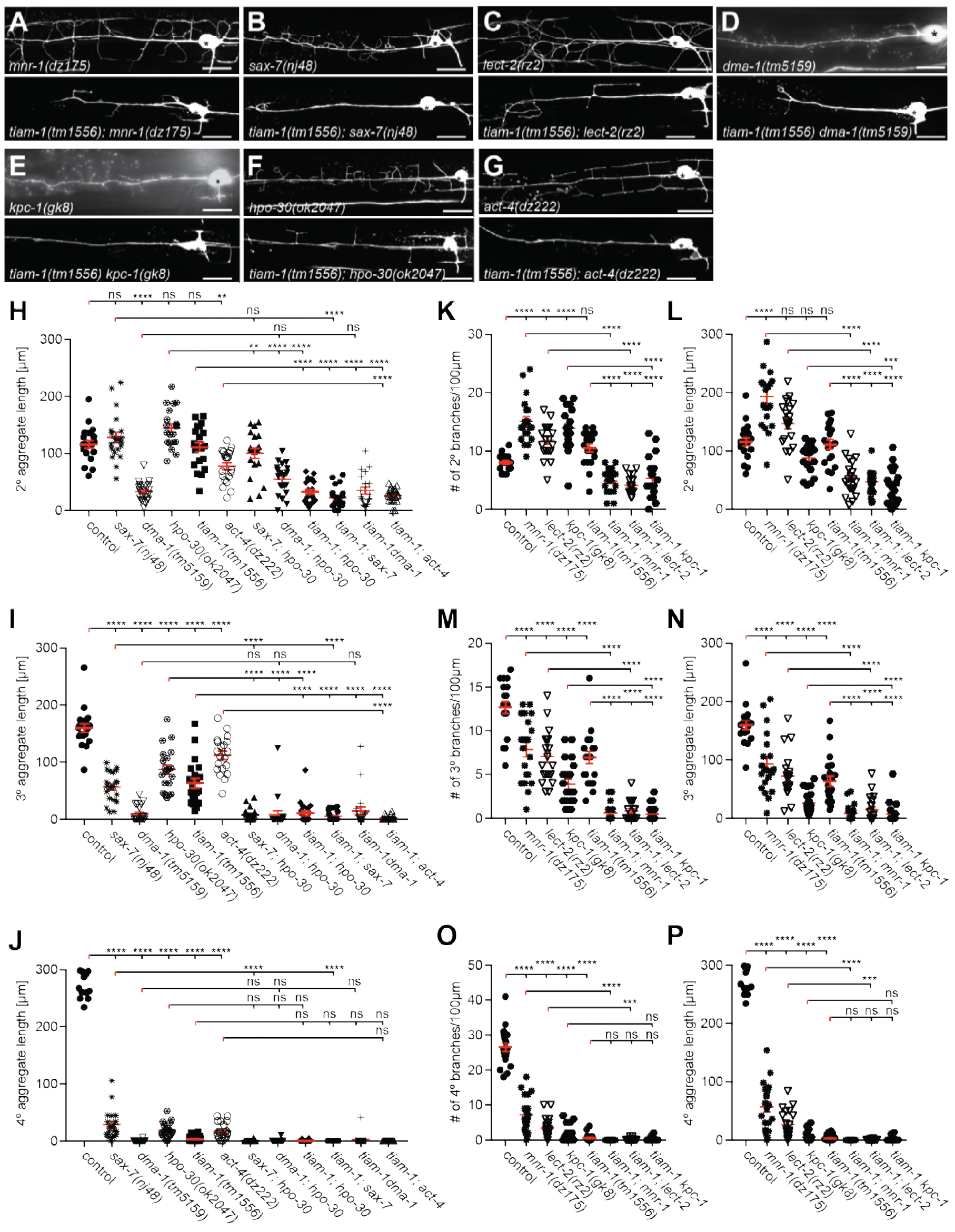
The Genetics of *hpo-30/Claudin, tiam-1/GEF*, and *act-4/Actin*. A. - G. Fluorescent images of animals expressing GFP in PVD neurons *(wdIs52 Is[PF49H12.4::GFP])* in the indicated genotypes. Anterior is to the left and scale bars indicate 20μm. H. - J. Quantification of the aggregate length of secondary, tertiary, and quaternary branches 100μm anterior to the PVD cell body. Data are represented as mean ± SEM. Statistical comparisons were performed using one-sided ANOVA with the Tukey correction. Statistical significance is indicated (ns: not significant, **p<0.05, ***p<0.005, ****p < 0.0005). n = 20 for all samples. K. - P. Quantification of the number (K,M,O) or aggregate length (L,N,P) of secondary, tertiary, and quaternary branches 100μm anterior to the PVD cell body. Some data in (K,M,O) is identical to data in Figure 2 and shown for comparison only. Data are represented as mean ± SEM. Statistical comparisons were performed using one-sided ANOVA with the Tukey correction. Statistical significance is indicated (ns: not significant, **p<0.05, ***p<0.005, ****p < 0.0005). n = 20 for all samples.

**Figure S4.**
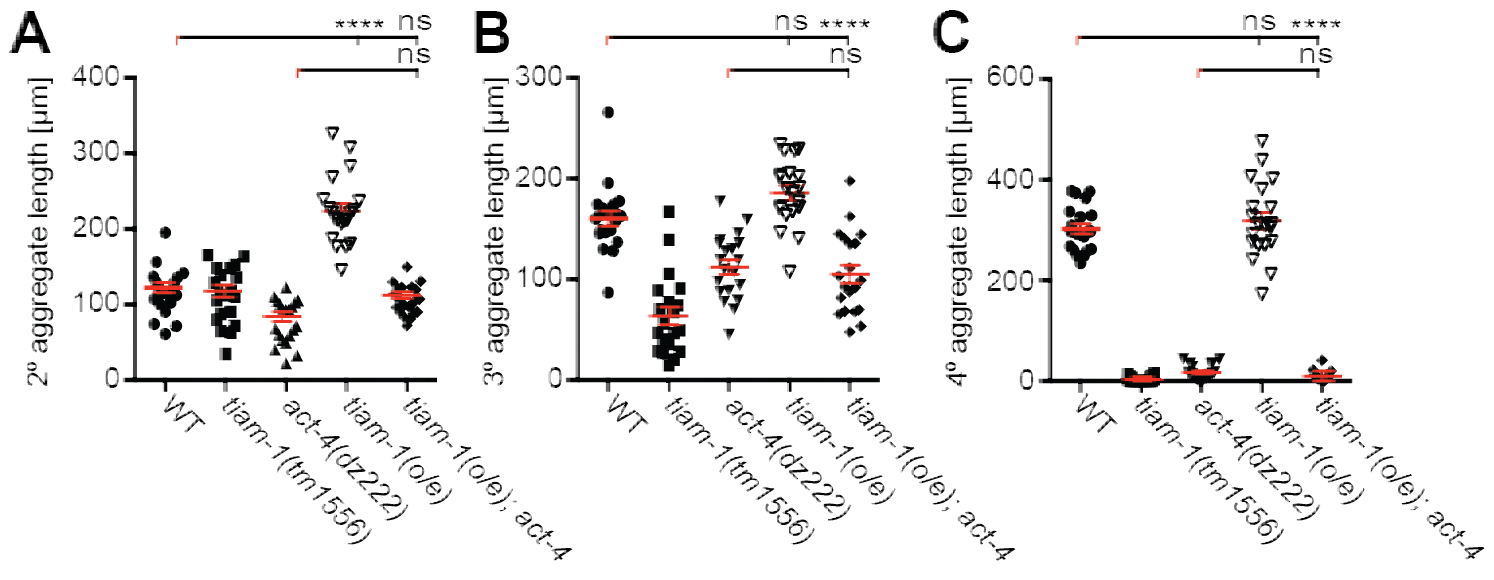
The *tiam-1/GEF* and *act-4/Actin* act downstream of the *dma-1/LRR-TM* receptor in PVD dendrites. Quantification of the aggregate length of secondary, tertiary, and quaternary branches 100μm anterior to the PVD cell body in the genotypes indicated. Data are represented as mean ± SEM. Statistical comparisons were performed using one-sided ANOVA with the Tukey correction. Statistical significance is indicated (ns: not significant, **p<0.05, ***p<0.005, ****p < 0.0005). n=20 for all samples.

**Figure S5.**
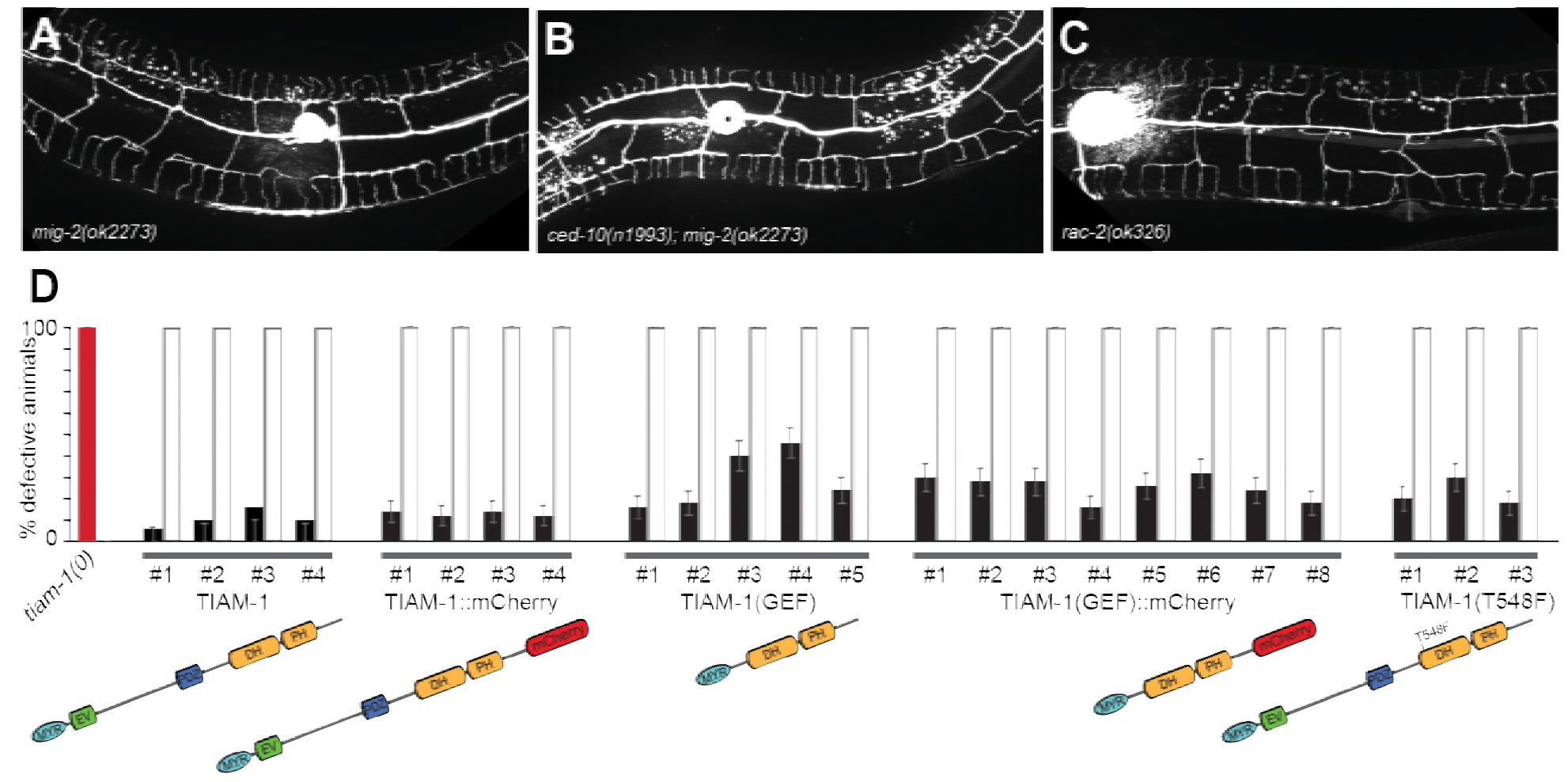
TIAM-1/GEF functions independently of GEF enzymatic activity. A. - C. Fluorescent images of animals in the indicated genetic backgrounds carrying a PVD cytoplasmic GFP reporter *(wdIs52)*. D. Quantification of animals with a defective PVD dendrite in transgenic rescue experiments. A red bar indicates the *tiam-1(tm1556)* mutant phenotype. Different transgenic lines (numbered #1-#N) are shown for each construct. Black bars indicate transgenic animals and white bars corresponding non-transgenic siblings. Shown is the percentage of animals with defective PVD dendritic arbors +/− the standard error of proportion.

**Figure S6.**
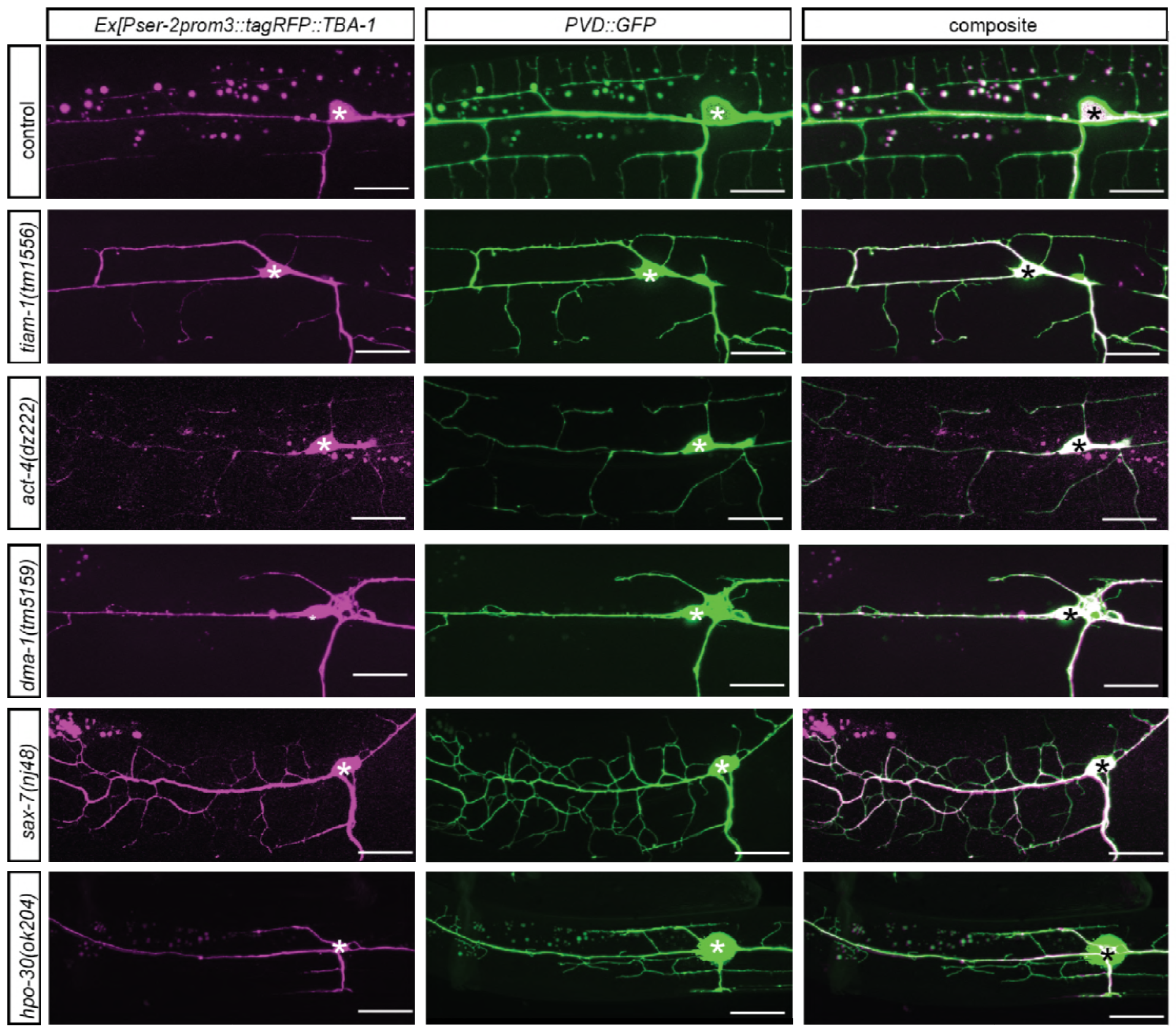
Localization of tagRFP::TBA-1 in different genetic backgrounds. Fluorescent images of animals in different genetic backgrounds carrying a microtubule reporter (*dzEx1569, Ex[Pser-2prom3::tagRFP::TBA-1]*, left panels), a PVD cytoplasmic GFP reporter (middle panels), and merged images (right panel). Genotypes are indicated on the left.

**Figure S7.**
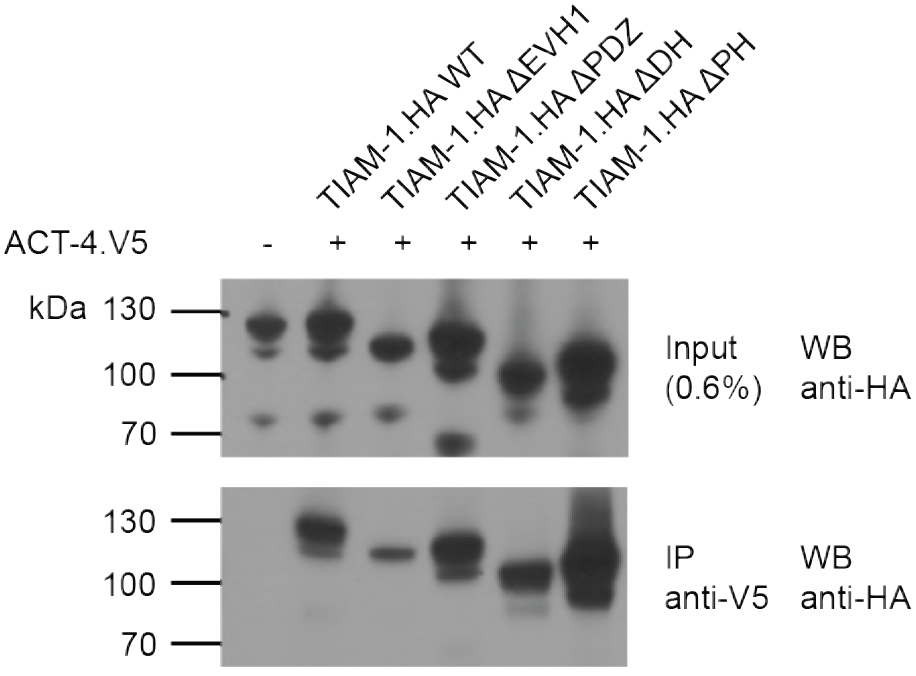
The DMA-1/LRR-TM, TIAM-1/GEF and ACT-4/Actin are part of the same biochemical complex. Western Blots of co-immunoprecipitation experiments. Transfected constructs are indicated above the panels. Antibodies used for immunopreciptation (IP) and Western Blotting (WB) are indicated. A molecular marker is on the left.

